# Hidden Spirals Underlie Cortical Traveling Waves

**DOI:** 10.1101/2025.11.03.686225

**Authors:** Anup Das, Jiawen Zhang, Erfan Zabeh, Luca Kolibius, Uma Mohan, Bard Ermentrout, Joshua Jacobs

## Abstract

Neural traveling waves exhibit diverse spatial patterns, but the neural dynamics that generate them remain unknown. Here, we asked whether the apparent diversity of traveling waves reflects distinct dynamical processes or a common underlying organization obscured in recorded brain activity. We analyzed intracranial recordings from humans performing a working memory task and developed FOCUS (**F**requency-flattened **O**scillators in **C**ortex **U**nmask **S**pirals), a coupled phase oscillator framework that accounts for spatial heterogeneity in local frequencies to uncover latent dynamics underlying the observed waves. FOCUS revealed covert spiral dynamics underlying traveling waves whose recorded spatial patterns showed no overt spiral organization. These hidden spirals emerged only after accounting for heterogeneous local frequencies, exposing dynamical structure that was inaccessible from the recorded wave patterns alone. The centers of the hidden spirals shifted across the cortex between memory encoding and retrieval, linking their spatial organization to distinct memory states. Together, these findings reveal that apparently complex spatial patterns of traveling waves can conceal latent spiral attractors, uncovering a previously inaccessible layer of neural organization in human memory.

## Introduction

Neural traveling waves are patterns of brain oscillations that progressively propagate across the cortex in specific directions to coordinate neural activity in space and time to support behavior (Mohan et al., 2024). Traveling waves propagate across the brain in a variety of spatial patterns (Das et al., 2026). Because propagating oscillations correlate with underlying neuronal activity (Jacobs et al., 2007), the spatiotemporal organization of traveling waves at each moment may indicate which brain areas are active and how they correspond to particular memory representations (Pinotsis et al., 2023; Pinotsis & Miller, 2023). However, despite their functional importance, the mechanisms that generate the diverse traveling wave patterns observed in brain activity remain elusive.

Our previous work (Das et al., 2026) used independent component analysis (ICA) to find a variety of spatial wave patterns in humans including plane waves, sources and sinks, and rotating waves. In addition, there were a large fraction of waves that were *complex wave* patterns that could not be classified as any of these simpler patterns. Complex waves are not simply a superposition of the simpler patterns as ICA would have detected this possibility. In this paper, we propose a novel mechanism for the formation of complex traveling waves (Agarwal et al., 2014; Bhattacharya et al., 2022; Das et al., 2026).

Previous work has suggested that spatial patterns of phases can be modeled as a locally connected network of phase oscillators; such models have been used to model synchronization, waves, and other spatial patterns in oscillatory systems (Abrams & Strogatz, 2004; Cohen et al., 1982; Dörfler & Bullo, 2014; Ermentrout et al., 2019; Ermentrout & Kopell, 1984; Kuramoto, 1984; Paullet & Ermentrout, 1994). For example, humans and animals exhibit plane waves, which propagate in particular directions related to current task processing. The simplest mechanism that can produce plane waves and concentric (sources and sinks) waves in a system of coupled phase are small differences in the local frequency. These plane waves originate from frequency gradients where faster oscillators lead slower ones (Ermentrout & Kopell, 1984; Zhang et al., 2018), creating a continuous spatial phase lag that manifests as a plane wave moving across the cortex. Similarly, sources (sinks) arise in two dimensions from a localized increase (decrease) in frequency at a single point, with the frequency gradually decreasing (increasing) with distance from that point (Hagan, 1981).

In addition, there is some evidence for traveling waves with other shapes, such as rotating waves or spirals (Das et al., 2026; Prechtl et al., 1997), although they are less common than plane waves (Das et al., 2026). Theoretical models show a strong basis for the existence for spiral/rotating waves. Rotating waves and synchrony occur in locally coupled identical phase oscillator models as phase-locked states (Bramburger, 2019; Paullet & Ermentrout, 1994; Shima & Kuramoto, 2004). Since rotating or spiral waves and synchrony are stable locked states to systems of identical oscillators, they are intrinsic to the network and do not require any external heterogeneities (Ding & Ermentrout, 2022; Paullet & Ermentrout, 1994; Shima & Kuramoto, 2004). In contrast, plane waves and sources/sinks originate from differences in local frequencies of the coupled oscillators. Because external inputs to oscillators typically affect their local frequency, we might consider any local differences in oscillator frequencies as arising from heterogenous external inputs.

Given their strong theoretical basis (Ding & Ermentrout, 2022; Paullet & Ermentrout, 1994), it might be considered surprising that, in practice, rotating waves such as spirals are observed less frequently than other wave types. Since frequency heterogeneities can give rise to plane or concentric waves, rotational dynamics may not be apparent in the recorded activity when frequency heterogeneities are present. This raises the possibility that some complex traveling wave patterns contain overt rotational or spiral morphology that are obscured by frequency heterogeneity. Therefore, we hypothesized that complex wave patterns are a consequence of heterogenous perturbations to networks of oscillators that have intrinsic spiral wave dynamics.

We tested this hypothesis by introducing FOCUS (**F**requency-flattened **O**scillators in **C**ortex **U**nmask **S**pirals), a novel computational technique in which we first estimate the frequencies and coupling in the coupled oscillator model to exactly fit the observed complex pattern. Then we gradually remove the frequency differences to reveal the *hidden* underlying intrinsic spiral dynamics. We applied this method to direct brain recordings from neurosurgical patients performing a working memory task. This computational method identified new spirals that were hidden within complex traveling wave patterns and were empirically invisible. These hidden spirals had novel features, such as showing that their center locations shifted across the cortex to distinguish different behavioral states, such as memory encoding, maintenance, and retrieval. Our findings reveal that the apparent complexity of traveling waves can conceal hidden spiral dynamics, exposing a previously inaccessible layer of neural organization during memory.

## Results

### Characterizing diverse spatial propagation patterns of traveling waves

To determine the generative mechanisms underlying the diverse traveling wave patterns observed in human brain activity, we first characterized their spatial and temporal organization in electrocorticographic (ECoG) recordings from five patients with implanted electrode grids as they performed a working memory task (**Methods, Figure 1, Supplementary Figure 1**). Our central goal was to design a novel computational framework based on coupled oscillator modelling to find hidden spirals withing these diverse spatial patterns, that appeared after removing plane and other traveling waves generated by frequency gradients. Therefore, we first used an analytical framework to characterize the observed spatial propagation patterns of traveling waves in the ECoG recordings, prior to investigating their underlying dynamics to find potential hidden signals.

**Figure 1:**
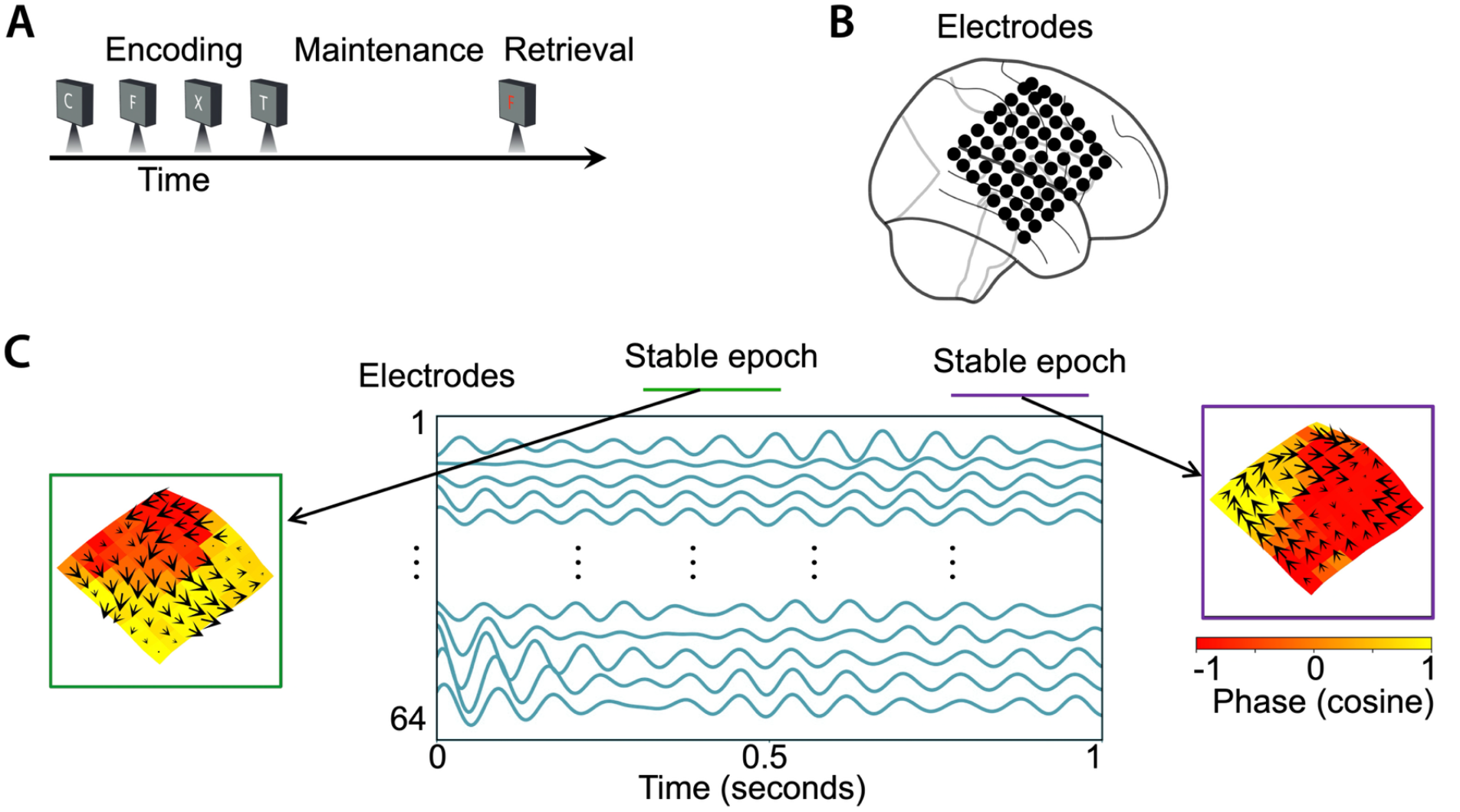
Working memory task and stable periods of traveling waves propagation. **(A) Sternberg working memory task.** Patients #1-5 (**Supplementary** Figure 1) performed multiple trials of a Sternberg working memory task where they were presented with a list of English letters to silently hold the identity of each item in memory and during the response period, indicated whether the probe was present in the just-presented list (**Methods**). **(B) An 8×8 ECoG grid in Patient #2. (C) Filtered signals and stable epochs.** We filtered the signals of the electrodes in each cluster at their peak frequencies (**Methods**). Shown here are the filtered signals from an example trial for the electrodes 1-64 in the ECoG grid shown in **B**. Colored horizontal lines (green and purple) denote spatiotemporally stable periods of wave propagation. Visual observation of the propagation of the absolute phase (Hilbert-transformed phase) across time showed stable periods of traveling waves. We used a localized circular-linear regression approach to estimate traveling waves in each patient individually and then identified stable periods of wave propagation (**Methods**). The traveling waves operated in the stable regime for a few tens of milliseconds, then they entered into the unstable regime where the stable wave pattern broke down and a new wave pattern emerged, and then finally moving onto a new stable regime. In the example shown here, the first stable epoch shows a counter-clockwise spiral wave, whereas the second stable epoch shows a clockwise spiral wave. Arrows denote the direction of the wave, lengths of the arrows denote wave strength, and colors denote the cosine of the phase.

We identified narrowband oscillations across cortical electrode clusters and quantified their instantaneous phase and spatial propagation using a windowed circular–linear regression framework (**Methods**). We extracted the direction and speed of instantaneous wave propagation at each electrode in an oscillation cluster. With this method, visual inspection of the phase patterns showed clear traveling waves, with oscillations advancing in consistent spatial patterns between neighboring electrodes (**Figure 1C**). We observed traveling waves across a range of oscillation frequencies spanning 6–20 Hz, with approximately 80% of electrodes exhibiting narrowband oscillatory activity in at least one frequency band. Because traveling waves can occur transiently (Das et al., 2026), we focused subsequent analyses on stable epochs in which their spatial organization remained stable over time (all *p’s* < 0.001, **Methods, Figure 1C**).

The stable traveling wave epochs exhibited diverse spatial patterns across subjects and memory states (**Figures 2, 3**), including plane, rotational, concentric (source/sink), and complex propagation (**Figure 2**). Individual cortical clusters frequently transitioned between these patterns across epochs, indicating that traveling wave activity was not restricted to a single spatial organization. To quantify this diversity over time, we used a statistical signal processing algorithm based on independent component analysis (ICA) (Fu et al., 2015; Li & Adalı, 2010), identifying recurring spatial propagation patterns, or *modes*, within each cortical cluster (**Figure 2, Methods**). Across subjects, individual clusters expressed a mean of nine significant modes, encompassing distinct combinations of plane, rotational, concentric (expanding or contracting), and complex wave patterns. We quantified the strength of each mode in an epoch by measuring the mode’s *activation function*, which measures the instantaneous magnitude and direction of the spatial traveling wave pattern represented by each mode. The results of this ICA procedure revealed a range of spatial traveling wave patterns in the individual subjects. For example, **Figure 2** shows that during the retrieval periods in Subject 2, the most dominant spatial pattern was a plane wave, explaining ∼ 40% of variance in the data, alongside less dominant rotational and complex spatial patterns of traveling waves.

**Figure 2:**
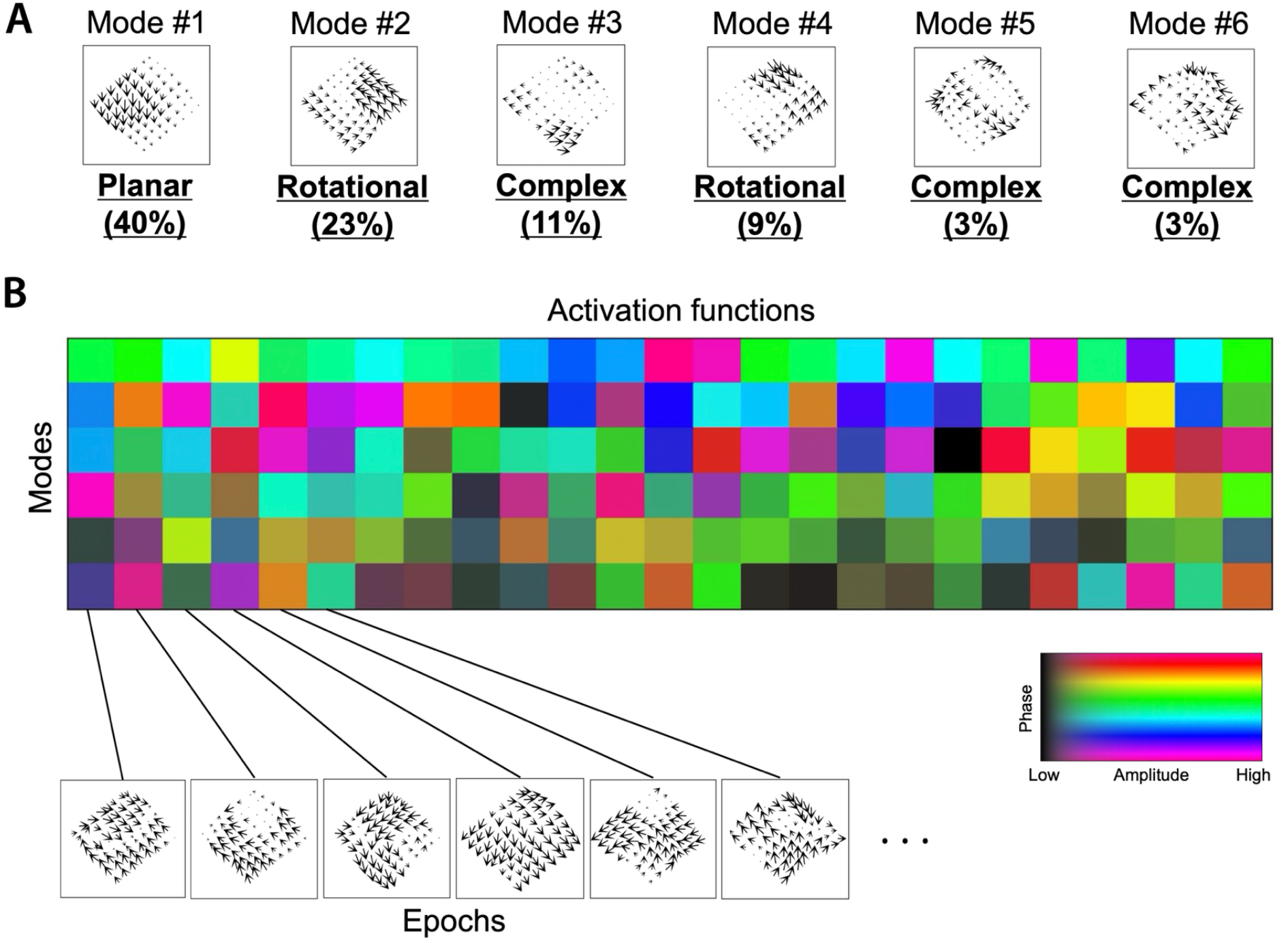
Independent component analysis (ICA) of traveling waves. Wave patterns across stable epochs are concatenated and passed as input to ICA **(Methods). (A, B) We extracted the modes and the independent activation functions (or, weights) as the outputs from the ICA (Methods).** Shown is an example of the application of the ICA method on the stable epochs during the retrieval period of the working memory task from Subject #2 (∼12 Hz traveling wave). Wave type and variance explained for each mode are indicated in **A**. We classified each of the modes as one of “planar”, “rotational”, “concentric” (“expanding” or “contracting”), or “complex” categories (**Methods**). Activation functions are complex numbers and shown in color with colorbar in **B**. Modes combine with the activation functions to form the stable epochs.

We next probed the potential functional role of different types of traveling waves, using multivariate analysis of variance (MANOVA) (**Methods**) to test for changes in activation functions between task conditions. Although parts of this approach had been employed previously (Das et al., 2026), an underappreciated aspect of the results was the presence of spiral waves in the human cortex. By testing for changes in activation functions over time, we found that traveling waves significantly shifted their behavior between stages of working memory processing (**Figure 3**). As an example, **Figure 3A** shows a cluster of electrodes with three distinct modes of traveling waves that shifted propagation direction between the stages of working memory. In this cluster, the plane wave pattern (mode 1) was nearly absent during encoding (as indicated by its low magnitude), relatively strongly present during maintenance, and was the strongest during the retrieval period (*p* < 0.001, MANOVA). Similarly, this cluster’s mode 2 showed a spiral wave during encoding and retrieval, but a source wave during the maintenance period (*p* < 0.001). This cluster’s mode 3 showed a complex spatial pattern of traveling waves that was the strongest during maintenance, and the weakest during encoding (*p* < 0.001). These state-specific traveling waves were widespread, as all five subjects had oscillation clusters showing significant shifts in direction or strength between task stages (all *p’s* < 0.05; also see **Figure 3B**).

**Figure 3:**
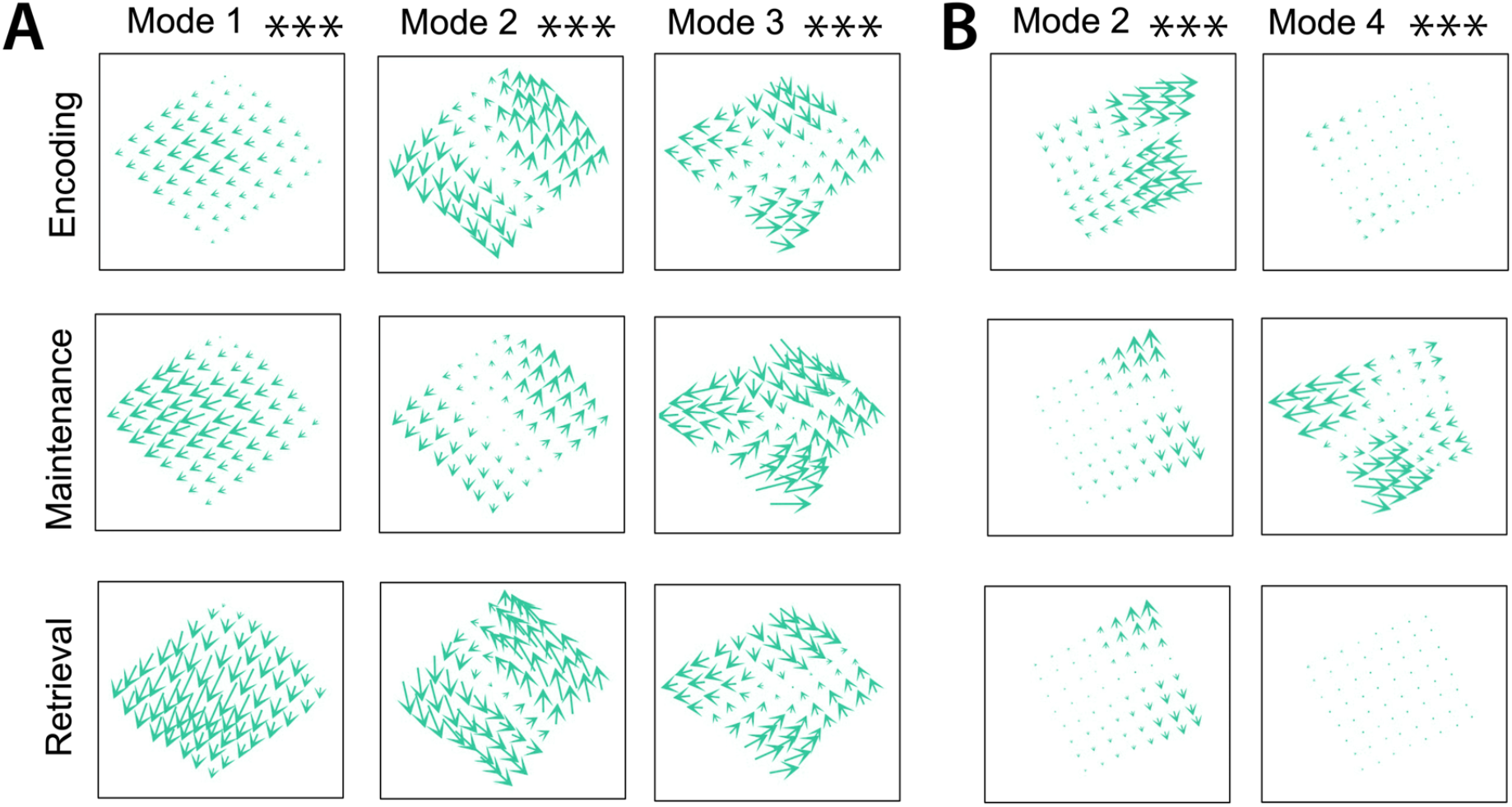
Traveling waves can distinguish behavioral states in human working memory. **(A) Top 3 mean modes of Subject #2 (∼ 12 Hz traveling wave) for the encoding, maintenance, and retrieval periods.** Traveling waves changed their direction and/or strength to distinguish different behavior in the working memory task. Encoding (N=1513), Maintenance (N=891), Retrieval (N=526). **(B) Similar to A, but for Subject #3.** This example shows two example mean modes (2 and 4) for the different task periods (∼ 7 Hz traveling wave) for Subject #3. Encoding (N=3294), Maintenance (N=2068), Retrieval (N=1111). (*** p < 0.001, MANOVA, FDR-corrected).

### Computational modeling of stable epochs using coupled phase oscillators

The diverse and complex traveling wave patterns observed in the ECoG recordings raised a central question: what underlying dynamics generate these distinct spatial expressions? Notably, complex wave patterns were more frequent than overtly rotational patterns (**Figures 2**, **3**). Because theoretical models show a strong basis for the existence for spiral/rotating waves (Ding & Ermentrout, 2022; Paullet & Ermentrout, 1994) and also because spatial frequency heterogeneity can shape the propagation of coupled oscillators, we reasoned that rotational dynamics could be obscured when multiple spatial frequency gradients contribute to the recorded wave pattern. Thus, the absence of an overt spiral morphology does not necessarily imply the absence of underlying rotational dynamics.

To test this possibility, we developed FOCUS, a coupled phase oscillator framework that accounts for the local frequencies and local interactions of neural populations. Coupled phase oscillators were shown to effectively model oscillations propagating across areas of cortex (Ermentrout & Kleinfeld, 2001). Previous research showed that two-dimensional arrays of nearest-neighbor coupled Kuramoto phase oscillators (Kuramoto, 1984) can model and produce stable patterns of rotating traveling waves (Ding & Ermentrout, 2022; Koller et al., 2024; Paullet & Ermentrout, 1994) (**Methods**). Since rotating or spiral waves and synchrony are stable locked states to systems of identical oscillators, they are intrinsic to the network and do not require any external heterogeneities (Ding & Ermentrout, 2022; Paullet & Ermentrout, 1994; Shima & Kuramoto, 2004). We therefore modeled each electrode as a Kuramoto phase oscillator coupled to its nearest neighbors and has its own oscillation frequency, and fit the model to the experimentally measured phase dynamics during stable traveling wave epochs (**Methods, Figure 4**). Frequencies of the Kuramoto oscillators can be thought of as perturbations on top of the narrowband frequency peaks that we previously detected. This computational framework therefore allowed us to separate the contribution of spatial frequency heterogeneity from the collective phase dynamics and determine whether rotational organization emerged in the resulting dynamical system.

**Figure 4:**
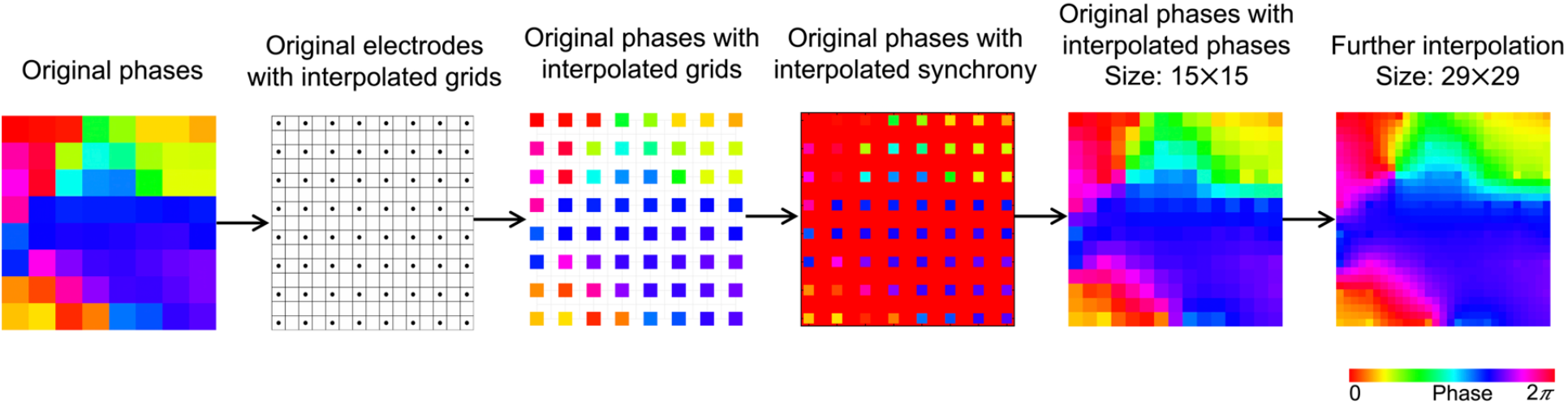
Interpolation procedure to reduce phase differences between adjacent oscillators in the coupled phase oscillator modeling framework. We embedded the 8×8 experimental ECoG array of phases in a larger 29×29 network to create smaller phase differences using a sine interpolation procedure (**Methods**). We initialized the sine model using synchrony states (zero phase values) for the missing values on the 15×15 grid with 64 fixed phase values corresponding to the original electrodes. We then used this matrix as the initial matrix for the forward solution to obtain the interpolation result. We used the same procedure to further interpolate the 15×15 grid to a 29×29 grid. This procedure produces a smoothly interpolated phase field with reduced nearest-neighbor gaps, thus ensuring dynamical stability, whose equilibrium solution we use in our Kuramoto oscillator model. Control analysis showed that this interpolation step has minimal impact on our results (**Methods**).

Phase oscillator systems model the rate of change of phase for a given oscillator as a function of its frequency and its coupling with its nearest neighbors (**Methods**). Applying this model to our recordings, by using the phases extracted from the stable epochs, the solution of this model will reveal the frequencies of an oscillator for each electrode. Because the intracranial electrodes in our dataset are relatively widely spaced, the relative phase differences between their respective oscillators can be quite large, which increases the chance that the patterns will be dynamically unstable (**Methods**). Therefore, we embedded our experimentally measured phases in a larger network using interpolation to create smaller phase differences (**Figure 4, Supplementary Figure 2, Methods**). This procedure produces a smoothly interpolated phase field with reduced nearest-neighbor gaps, ensuring dynamical stability, whose equilibrium solution we use in our Kuramoto oscillator model. Control analysis showed that this interpolation step has minimal impact on our results (**Methods**).

### Frequency flattening reveals hidden spirals

We next asked whether the complex traveling wave patterns observed in the recordings could reflect underlying latent dynamics that are obscured by spatial frequency heterogeneity. In a coupled network of Kuramoto oscillators, if the frequencies of all oscillators are the same, then, *synchrony*, spiral, and rotating waves can be generated (Paullet & Ermentrout, 1994). In contrast, in coupled oscillator networks, differences in frequencies can generate plane and concentric waves and therefore distort the underlying hidden signals, and subsequently appear as heterogeneous, complex spatial wave patterns when there are frequency heterogeneities (Paullet & Ermentrout, 1994). We therefore developed a *frequency flattening* procedure that progressively reduces the inferred frequency heterogeneity while tracking the corresponding phase locked solutions of the fitted oscillator network (**Figures 5**, **6, Methods**). This procedure provides an estimate of the phase organization that emerges when the effects of spatial frequency heterogeneity are removed.

**Figure 5:**
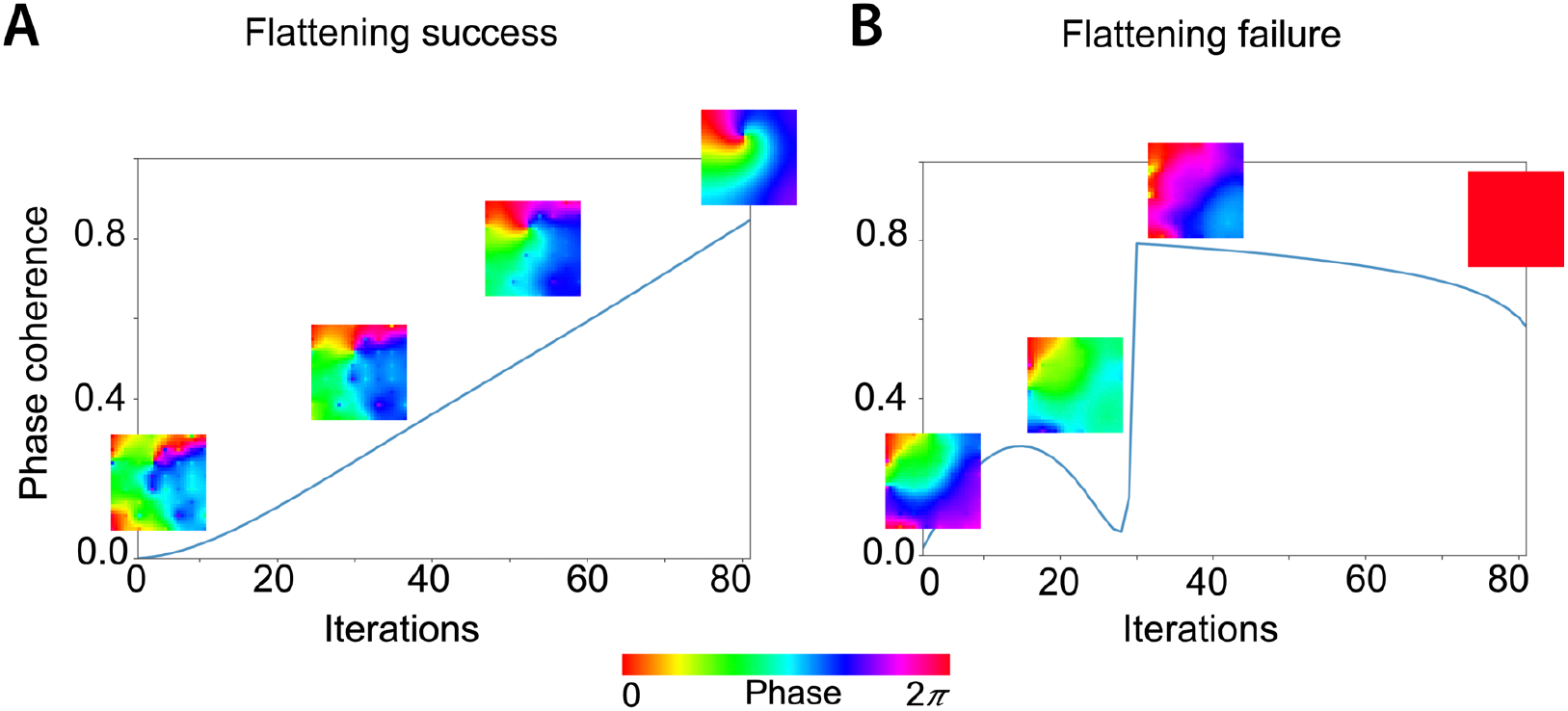
Order parameter trajectories illustrating one successful (A) and one unsuccessful (B) flattening process. The vertical axis represents: 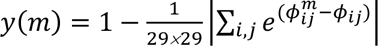, quantifying the phase coherence relative to the initial phase data pattern, where m denotes the iteration number. Flattening success involves a smooth order parameter trajectory, whereas a flattening failure involves a discontinuity (at m = 30 in **(B)**), highlighting a sharp phase transition occurring within a single forward iteration. Also shown in **(A)**, four example phase patterns for m = 1, 27, 54, and 81 respectively. Also shown in **(B)**, four example phase patterns for m = 1, 29, 30, and 81 respectively.

**Figure 6:**
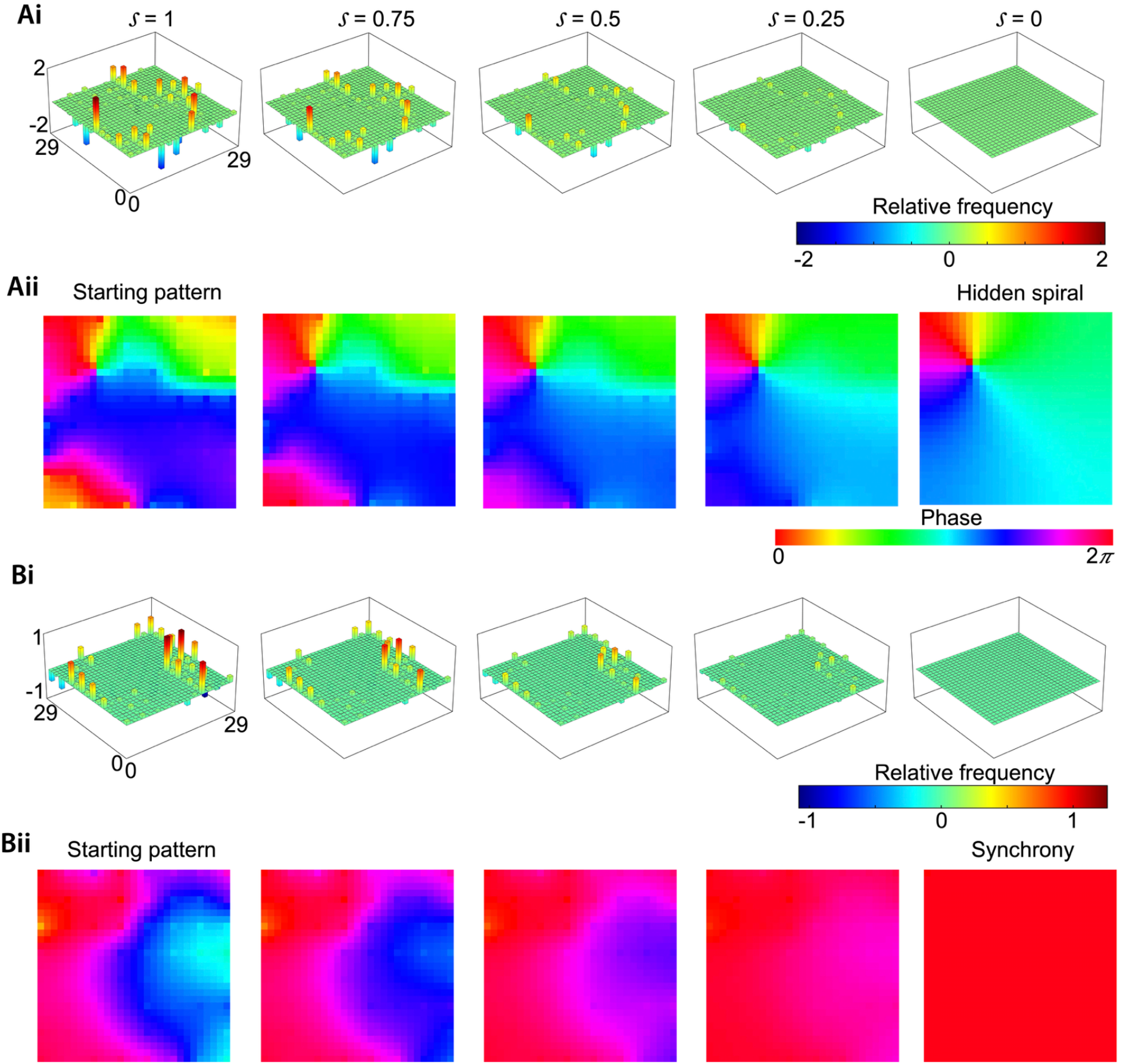
Frequency and phase propagation during the iterative frequency flattening procedure. **(Ai)** Iterative frequency flattening where the complex spatial phase pattern in **Aii** had a hidden spiral. Scales (1 − mℎ, where m = 0, 20, 40, 60, 80, see **Methods**) of this flattening are noted on top of each panel. **(Aii)** Corresponding phase propagation. **(Bi)** Iterative frequency flattening where the complex spatial phase pattern in **Bii** had a synchrony. **(Bii)** Corresponding phase propagation.

In this frequency flattening procedure, we first fit the Kuramoto model to the data. Then, by perturbing the model, we iteratively remove the plane and concentric traveling waves (frequency gradients). The steady-state solution of this computational framework after frequency flattening is the underlying *hidden* phase pattern. **Figure 6A** shows an example of this method where frequency flattening of a complex spatial wave pattern from an example stable epoch leads to a spiral. This spiral, which was obscured by frequency gradients, was visible once the frequency gradients that led to a complex spatial pattern were removed. **Figure 6B** shows an example of another complex wave pattern, from a different stable epoch where frequency flattening leads to synchrony.

The frequency flattening procedure is a method for estimating the phases in the network as the parameter of the network is shifted. In this case, the parameter that we are shifting is the phase coherence of the oscillator network (Krauskopf et al., 2007) (**Figure 5**). If the phase coherence increases smoothly without any jumps, then eventually the hidden phase pattern is revealed, and we call it *flattening success* (Krauskopf et al., 2007) (**Figure 5A**). By nature of the multi-stability of the solutions that this method can find, it is possible that flattening can result in multiple patterns of stable solutions and therefore, it can jump to different sets of phase-locked patterns (**Figure 5B**). For example, **Figure 5A** shows a smooth solution branch where a complex spatial pattern from an example stable epoch slowly converges to a spiral, which we refer to a *flattening success*. In contrast, **Figure 5B** shows a different solution branch, where flattening causes a complex spatial pattern to abruptly jump to a different phase-locked spatial pattern, eventually converging to a synchronous solution. We term this as a *flattening failure*. Therefore, to ensure that our results from frequency flattening are consistent and unique, we compare our results with the patterns that result from running a reversed version of the process. In this reverse process, we take the final flattened pattern and gradually decrease the phase coherence by gradually increasing the frequency heterogeneity by solving the same differential equation sequence in the opposite order. We then compare this converged phase pattern, where the frequency is at its original level, with the interpolated phase pattern we started with. If those phase networks match, we conclude that the results are unique and consistent. Using this procedure of frequency flattening, we found that ∼ 23% of all stable epochs in our experimental data satisfied this criterion of being consistent and unique, with ∼ 10% hidden spirals and ∼ 13% synchrony. The percentage of hidden spirals was statistically significant compared to a surrogate distribution (*χ*^2^ test, *p* < 0.001, **Methods**).

### Hidden spirals are distinct from overt rotational waves

We next asked whether the hidden spirals identified by frequency flattening simply reflected rotational waves already present in the recorded activity. If so, the modeling framework would merely recover an overt spatial pattern rather than reveal previously inaccessible dynamics. We therefore directly compared the frequency-flattened spiral solutions with rotational waves detected in the corresponding stable epochs using the ICA-based statistical analysis (**Methods**).

Across subjects and memory states, only ∼10% of the hidden spirals overlapped with rotational waves identified by ICA in the original recordings. Thus, most hidden spirals emerged from epochs that did not exhibit a corresponding overt rotational pattern. This limited overlap indicates that the model-derived spirals cannot be explained simply by the presence of rotational waves in the recorded activity.

Instead, the analysis reveals a distinction between overt wave morphology and underlying dynamical organization: traveling waves that appear spatially complex in the recordings can exhibit latent spiral organization after accounting for spatial frequency heterogeneity. Hidden spirals therefore represent a dynamical structure that is largely not expressed in the observed spatial morphology of the traveling waves.

### Hidden spirals are diverse

Having established that hidden spirals are distinct from overt rotational waves, we next systematically quantified their features such as their orientations and locations, fitting models separately during encoding, maintenance, and retrieval (**Figure 7, Methods**).

**Figure 7:**
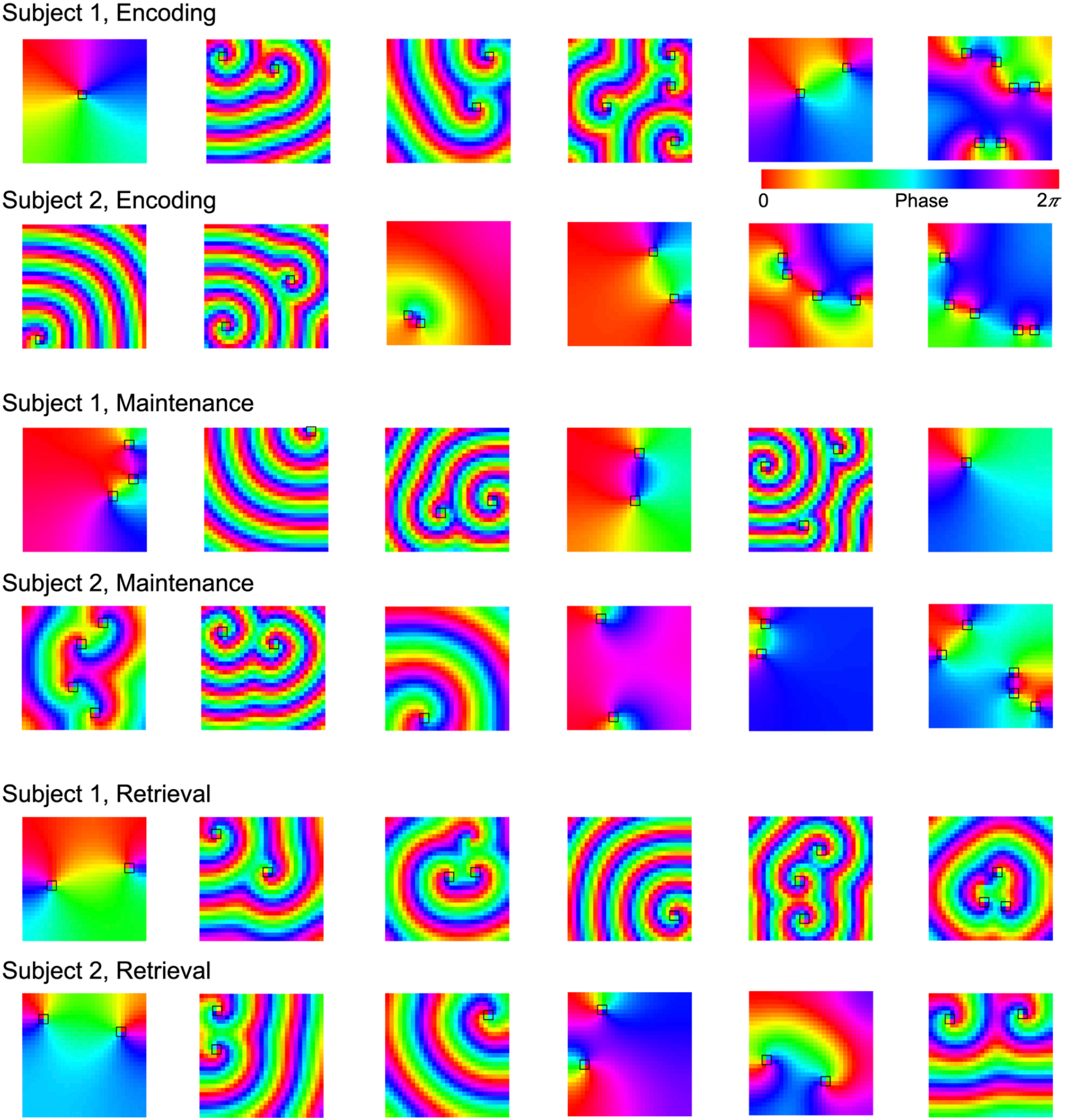
Hidden spirals are diverse. Shown here are 6 example hidden spirals for each task condition and for each Subject 1, 2. Black squares denote the core of the spirals. Each panel here corresponds to one stable epoch. Note that there can be multiple hidden spirals for the same stable epoch. See text for the description of these hidden spirals.

We first identified the location of the spiral core (black boxes in **Figure 7**) and from this, we identified the orientation (clockwise/counter-clockwise) of the hidden spiral (**Figure 8, Methods**). Hidden spirals exhibited both clockwise and counter-clockwise orientations, corresponding to temporal–frontal–parietal (TFP) and temporal–parietal–frontal (TPF) propagation, respectively, and both orientations occurred across subjects and task conditions (**Figure 7**). Their spatial organization was likewise heterogeneous: spiral cores occurred at different locations across the cortical electrode grids and varied across memory states (**Figures 11**, **12**).

**Figure 8:**
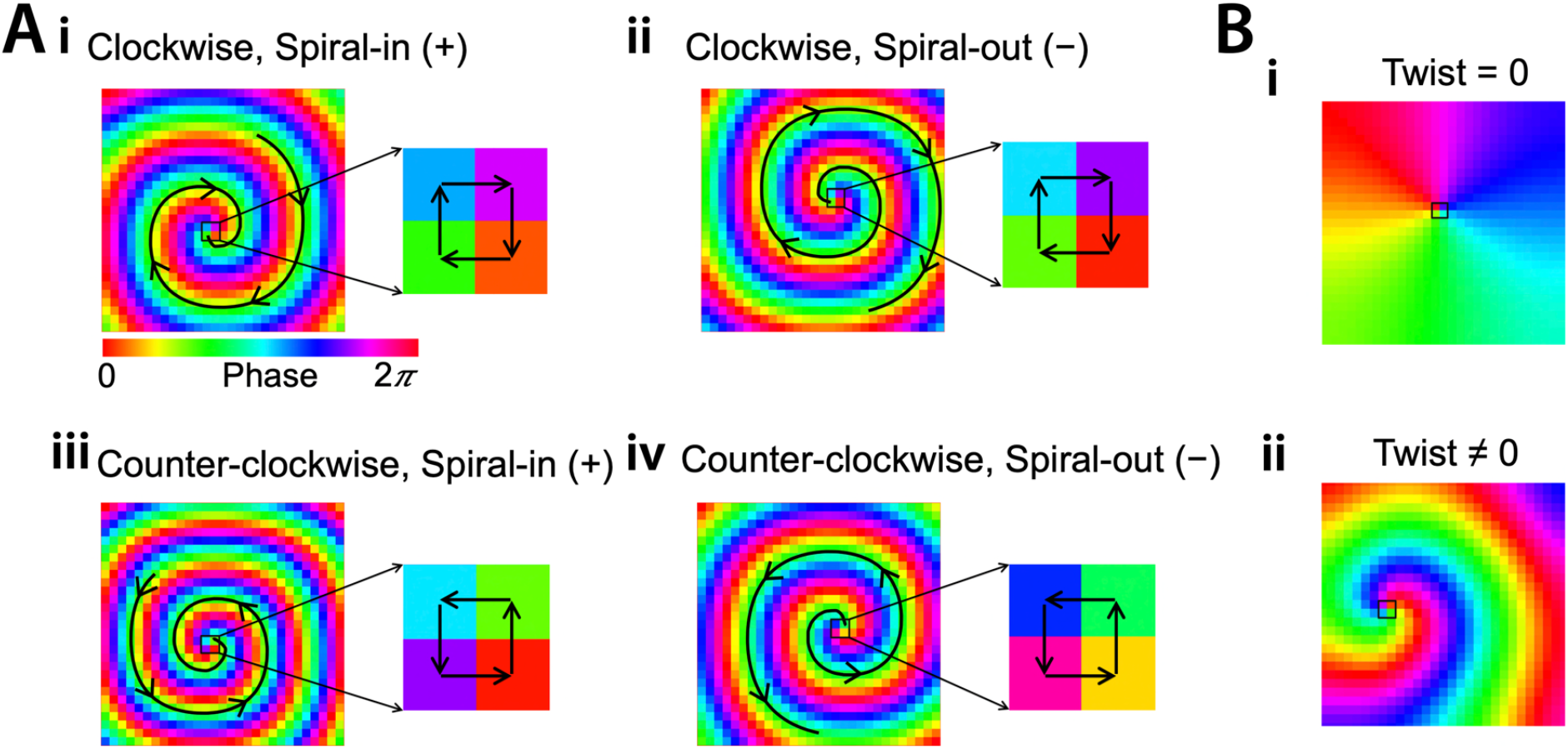
Orientation and twist of the hidden spirals. (A) Determining inward and outward spirals from the twist sign. The sign (positive(+)/negative(−)) of the twist metric T indicates whether it’s an inward (spiral-in) or outward (spiral-out) hidden spiral respectively (**Methods**). Twist of a hidden spiral was quantified using the metric 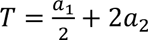, where a_1_and a_2_ are the coefficients of cos φ and cos 2φ in the coupling function respectively (**Methods**). **Ai** and **Aii** show two hidden spirals with the same orientation (clockwise or TFP), as indicated in the zoomed-in versions by the color change of the phases at the 2×2 core. However, they differ in their twist-orientation, i.e., **Ai** has an inward twist (spiral-in), as indicated by the positive sign of the twist, whereas **Aii** has an outward twist (spiral-out), as indicated by the negative sign of the twist. Iso-phase lines with arrows indicate the direction (spiral-in or spiral-out) of the twist. Similarly, **Aiii** and **Aiv** show two hidden spirals with the same orientation (counter-clockwise or TPF), as indicated in the zoomed-in versions by the color change of the phases at the 2×2 core. However, they differ in their twist-orientation, i.e., **Aiii** has an inward twist (spiral-in), as indicated by the positive sign of the twist, whereas **Aiv** has an outward twist (spiral-out), as indicated by the negative sign of the twist. Iso-phase lines with arrows indicate the direction (spiral-in or spiral-out) of the twist. Hidden spiral in **Ai** corresponds to the retrieval period in Subject 4. Hidden spiral in **Aii** corresponds to the encoding period in Subject 1. Hidden spiral in **Aiii** corresponds to the retrieval period in Subject 2. Hidden spiral in **Aiv** corresponds to the encoding period in Subject 3. **(B) Twist value of hidden spirals.** Higher the magnitude of the twist value, higher the twist of the hidden spiral. Shown are no-twist (**i**) and high-twist (**ii**) hidden spirals. Hidden spiral in **Bi** corresponds to the encoding period in Subject 1. Hidden spiral in **Bii** corresponds to the retrieval period in Subject 2.

Individual stable epochs could also contain multiple hidden spirals, sometimes with several cores distributed across the cortical grid and, in some cases, with different orientations within the same epoch (**Figure 7, Supplementary Figure 3**). The spatial frequency of the spirals also varied substantially across epochs. Thus, the latent rotational organization was itself dynamic and spatially diverse, rather than corresponding to a single fixed spiral configuration.

Together, these findings show that the dynamical structure underlying cortical traveling waves is flexible across memory states, with hidden spiral organization varying in location, orientation, multiplicity, and spatial scale. This state-dependent organization provides a potential dynamical substrate through which distributed cortical activity can be reorganized during different phases of memory processing.

### Same orientation hidden spirals can have distinct twists

In addition to characterizing the orientation of the hidden spirals, we also quantified the twists of the hidden spirals (**Methods, Figure 8**). We observed hidden spirals of varying degree of twists (**Figures 7**, **8B**). For example, during encoding, Subject 1 showed hidden spirals which had no twist (panel 1 in **Figure 7**). These no-twist spirals were also present during the maintenance period of Subject 1 (panel 6). We observed high-twist spirals across task conditions, for example, during the encoding period in Subject 2 (panel 1), during the maintenance period in Subject 1 (panel 2), and during the retrieval period in Subject 2 (panel 3).

An additional novel feature of the hidden spirals was that the twist-orientation of these hidden spirals could be distinct even if they had the same orientation (**Figure 8**). For example, **Figures 8Ai** and **8Aii** show two hidden spirals both of which have identical, clockwise (TFP) orientations as evident from the center of the spirals, however, their twist-orientations are different. **Figure 8Ai** shows a hidden spiral with a spiral-in twist, corresponding to the retrieval period in Subject 4, whereas **Figure 8Aii** shows a hidden spiral with a spiral-out twist, corresponding to the encoding period in Subject 1. Similarly, **Figures 8Aiii** and **8Aiv** show two hidden spirals both of which have counter-clockwise (TPF) orientations, however, their twist-orientations are different. One has a spiral-in twist (**Figure 8Aiii**), corresponding to the retrieval period in Subject 2, and the other has a spiral-out twist (**Figure 8Aiv**), corresponding to the encoding period in Subject 3.

### Hidden spirals are anticorrelated with plane waves

Inspecting our results, we observed that the latent dynamical organization of a traveling wave may depend on its observed spatial expression. Stable epochs with more complex spatial patterns more often contained a hidden spiral, compared to those with less complex spatial patterns, which generally showed only synchrony following frequency flattening (**Figure 6**). We therefore asked whether the presence of hidden spirals was systematically related to the empirically observed traveling wave patterns.

For each stable epoch in which frequency flattening converged, we quantified the relationship between the activation of each ICA-derived wave mode and the probability of a hidden spiral (**Figures 9**, **10, Methods**). In an example from Subject 2 during encoding, stronger plane wave activation was associated with a lower probability of a hidden spiral (Pearson *r* = −0.40, *p* < 0.001), whereas several complex wave modes showed positive associations with hidden spirals (Pearson *r* = 0.13, *p* < 0.001 for mode 4; Pearson *r* = 0.21, *p* < 0.001 for mode 5, **Figure 9D**). However, presence of a complex spatial shape did not always increase the probability of detecting a hidden spiral, for example, in mode 3, the presence of a complex spatial pattern was uncorrelated with the detection of a hidden spiral (Pearson *r* = 0.01, *p* = 0.844, **Figure 9D**). For this subject, during encoding, we also observed a significant positive correlation in mode 2, where the presence of a rotational wave increased the probability of detecting a hidden spiral (Pearson *r* = 0.12, *p* < 0.001, **Figure 9D**).

**Figure 9:**
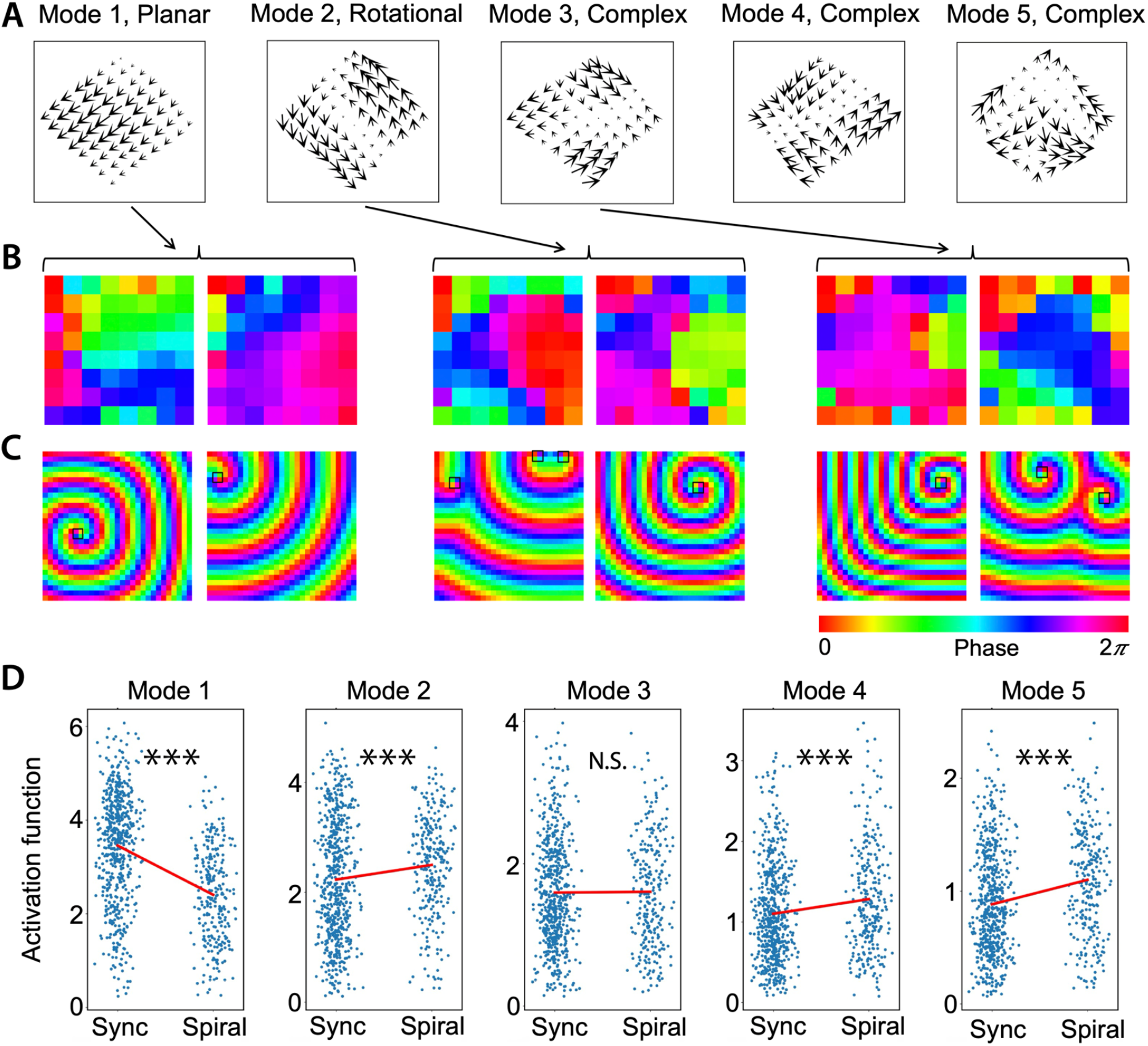
Hidden spirals are related to the ICA modes. **(A)** First five modes, with the wave types denoted on top, for the encoding period in Subject 2. Note the presence of multiple shapes of complex wave modes. **(B, C)** Single trial stable epoch phase examples in **B** corresponding to the different wave types in **A**. Hidden spirals are shown in **C** corresponding to the stable epochs in **B**. **(D)** Correlation between the activation functions from the ICA and the hidden pattern (synchrony/spiral) for each of the five modes shown in **A**. The corresponding modes are noted on top of each panel. N = 600 for synchrony and N = 275 for hidden spiral. (*** p < 0.001, N.S. Not significant, Pearson r, FDR-corrected)

**Figure 10:**
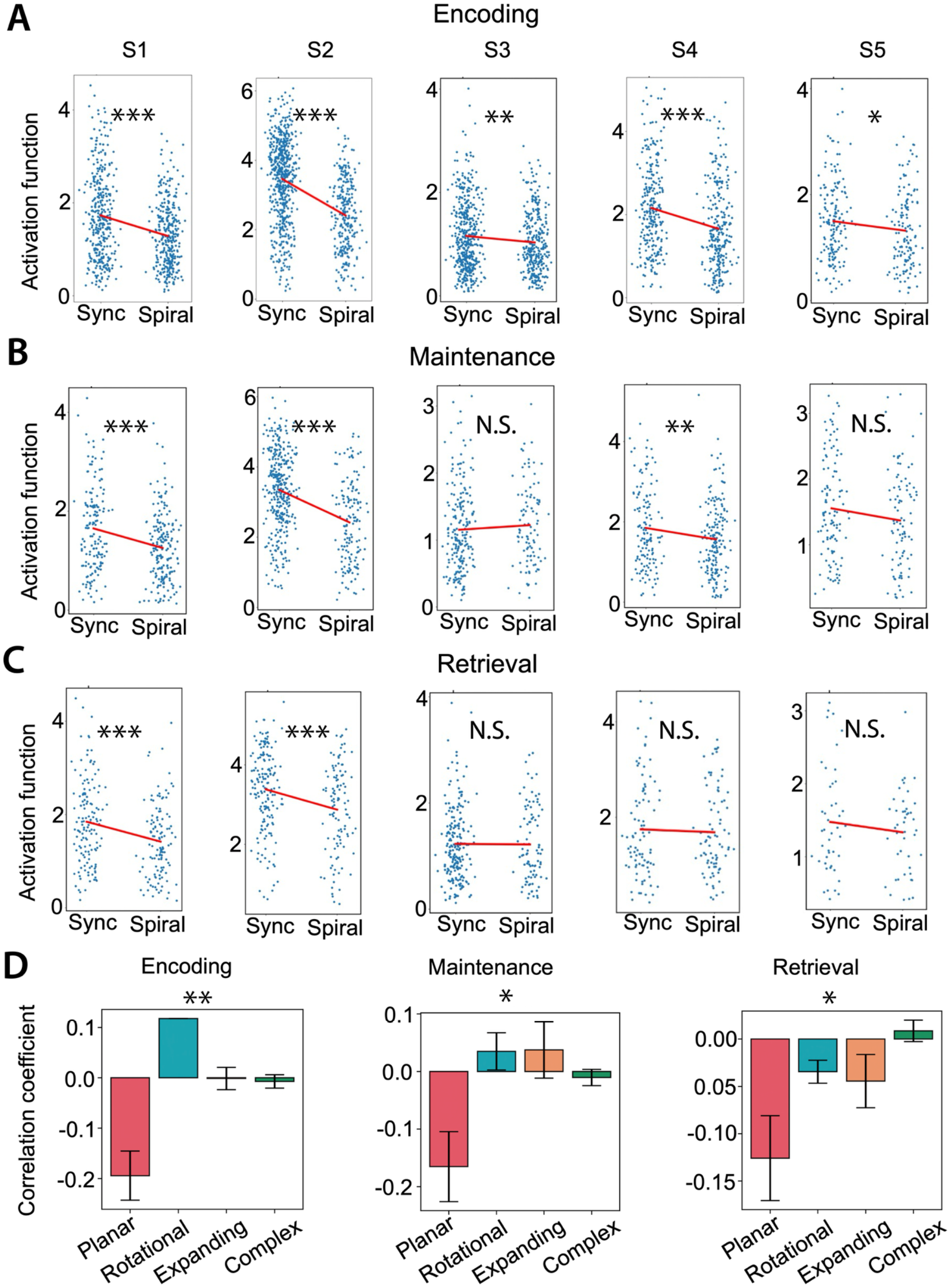
Hidden spirals are negatively correlated with the plane wave modes. Analyzing the correlation between the activation functions of the plane wave modes and the hidden pattern (synchrony/spiral) revealed that the plane wave modes are negatively correlated with the hidden spirals, a trend observed across all subjects and task conditions encoding **(A)**, maintenance **(B)**, and retrieval **(C)**. S1-S5 denote subjects 1-5. (*** p < 0.001, ** p < 0.01, * p < 0.05, N.S. Not significant, Pearson r, FDR-corrected). For encoding (synchrony), N = 352, 600, 479, 258, 168 for S1-S5 respectively. For encoding (hidden spiral), N = 334, 275, 326, 268, 140 for S1-S5 respectively. For maintenance (synchrony), N = 156, 354, 213, 138, 110 for S1-S5 respectively. For maintenance (hidden spiral), N = 196, 157, 100, 182, 93 for S1-S5 respectively. For retrieval (synchrony), N = 143, 171, 206, 92, 45 for S1-S5 respectively. For retrieval (hidden spiral), N = 124, 111, 93, 82, 45 for S1-S5 respectively. **(D)** Population level analysis (across all modes) revealed the same finding, where the probability of detecting a hidden spiral gets reduced for a planar (P) wave compared to other wave types such as rotational (R), expanding/contracting (E), and complex (C), for the encoding, maintenance, and retrieval task conditions. (** p < 0.01, * p < 0.05, Kruskal-Wallis, FDR-corrected). For encoding, N = 5, 1, 6, 32 for P, R, E, and C wave types respectively. For maintenance, N = 5, 4, 5, 34 for P, R, E, and C wave types respectively. For retrieval, N = 5, 3, 3, 38 for P, R, E, and C wave types respectively.

Across subjects and memory states, this relationship was remarkably consistent: plane wave activation was negatively correlated with hidden spiral occurrence in 10 of 15 plane modes and all five subjects, with significant negative correlations observed across encoding, maintenance, and retrieval (Pearson *r* ≤ −0.10, *p’s* < 0.05; **Figures 10A–C**). Moreover, the association with hidden spirals was significantly weaker for plane waves than for rotational, concentric (expanding/contracting), and complex wave patterns across all three memory states (encoding, *p* < 0.01; maintenance, *p* < 0.05; retrieval, *p* < 0.05, Kruskal–Wallis tests; **Figure 10D**).

This anticorrelation provides an important link between the observed morphology of traveling waves and their underlying dynamical organization. When plane wave expression was stronger, hidden spirals were less likely to be detected; conversely, epochs with stronger complex wave expression were more likely to contain hidden spirals. Thus, the emergence of latent rotational dynamics was not independent of the spatial structure expressed in the recordings.

These relationships are consistent with theoretical predictions that spatial heterogeneity in local frequencies can distort latent rotational dynamics into apparently spatially complex traveling wave patterns (Paullet & Ermentrout, 1994). Our findings extend this prediction to human cortical recordings: the prevalence of plane wave expression decreases as hidden spiral organization becomes more likely, whereas more heterogeneous wave morphologies are associated with the emergence of latent rotational dynamics. Together, these results provide a direct link between the diversity of observed traveling wave patterns and the latent dynamics that generate them. The spatial morphology seen in the recordings therefore carries information about, but does not fully reveal, the underlying dynamical state of the cortical network.

### Hidden spiral centers shift across the cortex between behavioral states

We next asked whether the latent spiral dynamics were reorganized across behavioral states of working memory. Specifically, we examined whether the spatial locations of TFP and TPF hidden spiral centers changed between encoding, maintenance, and retrieval, analyzing each subject separately (**Figure 11**).

**Figure 11:**
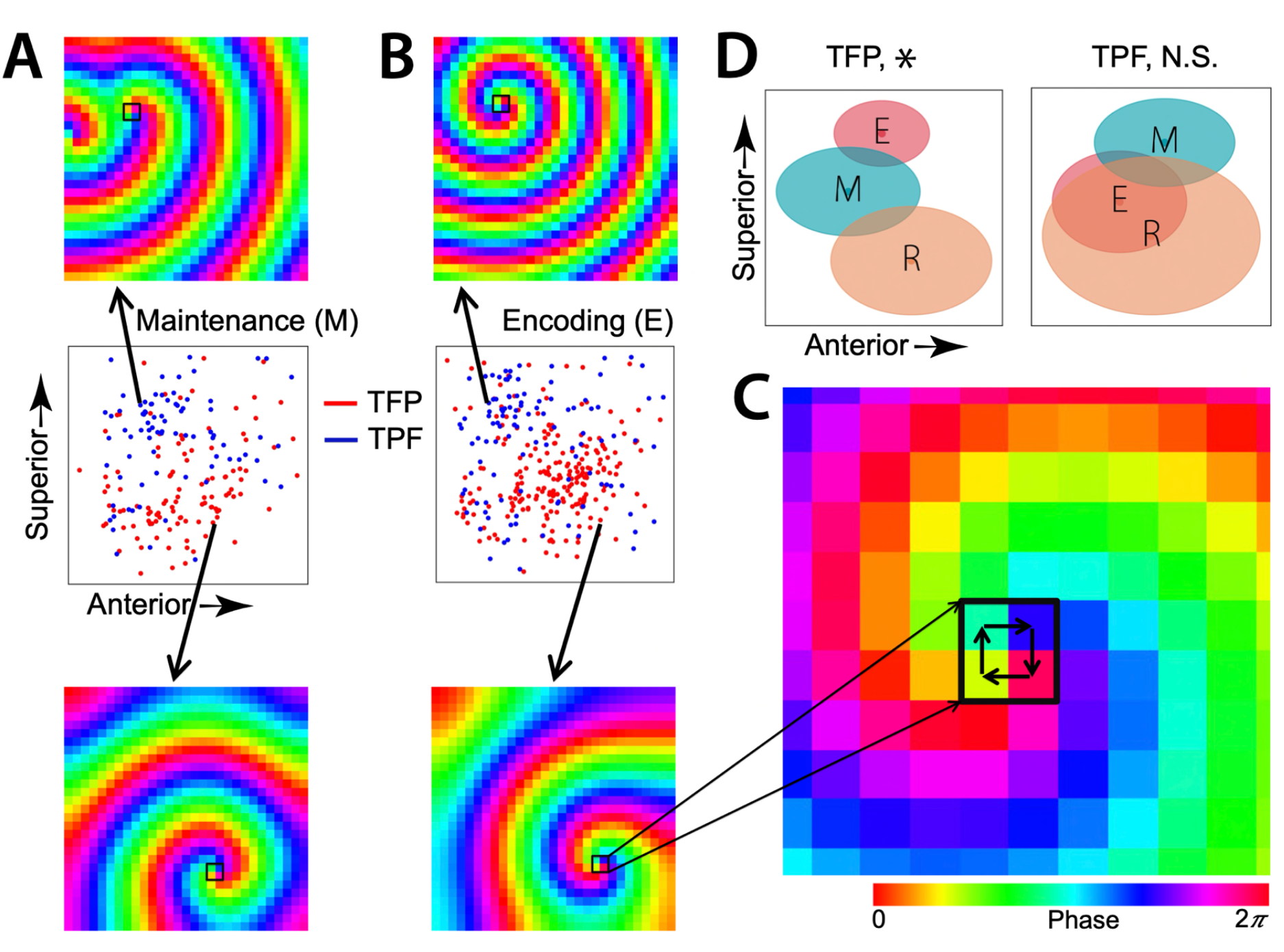
The center of the TFP and TPF spirals shift across the cortex to distinguish different behavioral states encoding (E), maintenance (M), and retrieval (R). **(A) Spatial distribution of the centers of the TFP (red) and TPF (blue) hidden spirals during the maintenance period in Subject 2.** Also shown are example TFP and TPF hidden spirals corresponding to the spatial location of their center. **(B) Same as in (A), but for the encoding period. (C) Shows a zoomed-in version of an example hidden spiral in (B).** Note the TFP orientation of the hidden spiral as indicated by the color change of the phases in a TFP orientation at its 2×2 core. **(D) TFP hidden spirals for Subject 2 shifted the location of their centers across the cortex to distinguish behavioral states, in contrast to the TPF hidden spirals where the shift in the spatial location across behaviors was not statistically significant.** For each ellipse (task period), the major axis (horizontal axis) denotes the standard-error-of-the-mean (SEM) for the Y-coordinate and the minor axis (vertical axis) denotes the SEM for the Z-coordinate, of the centers of the hidden spirals. (* p < 0.05, N.S. Not significant, MANOVA, FDR-corrected).

Visual inspection revealed clear state-dependent shifts in spiral-center locations across the cortical grids. To quantify these changes, we used multivariate analysis of variance (MANOVA; **Methods**) to test whether the two-dimensional positions of hidden spiral centers differed across task conditions. TFP hidden spirals showed significant spatial shifts across encoding, maintenance, and retrieval in all five subjects (all *p’s* < 0.05; **Figure 12A**), whereas TPF spirals showed significant shifts in three of five subjects (*p’s* < 0.05, **Figure 12B**).

**Figure 12:**
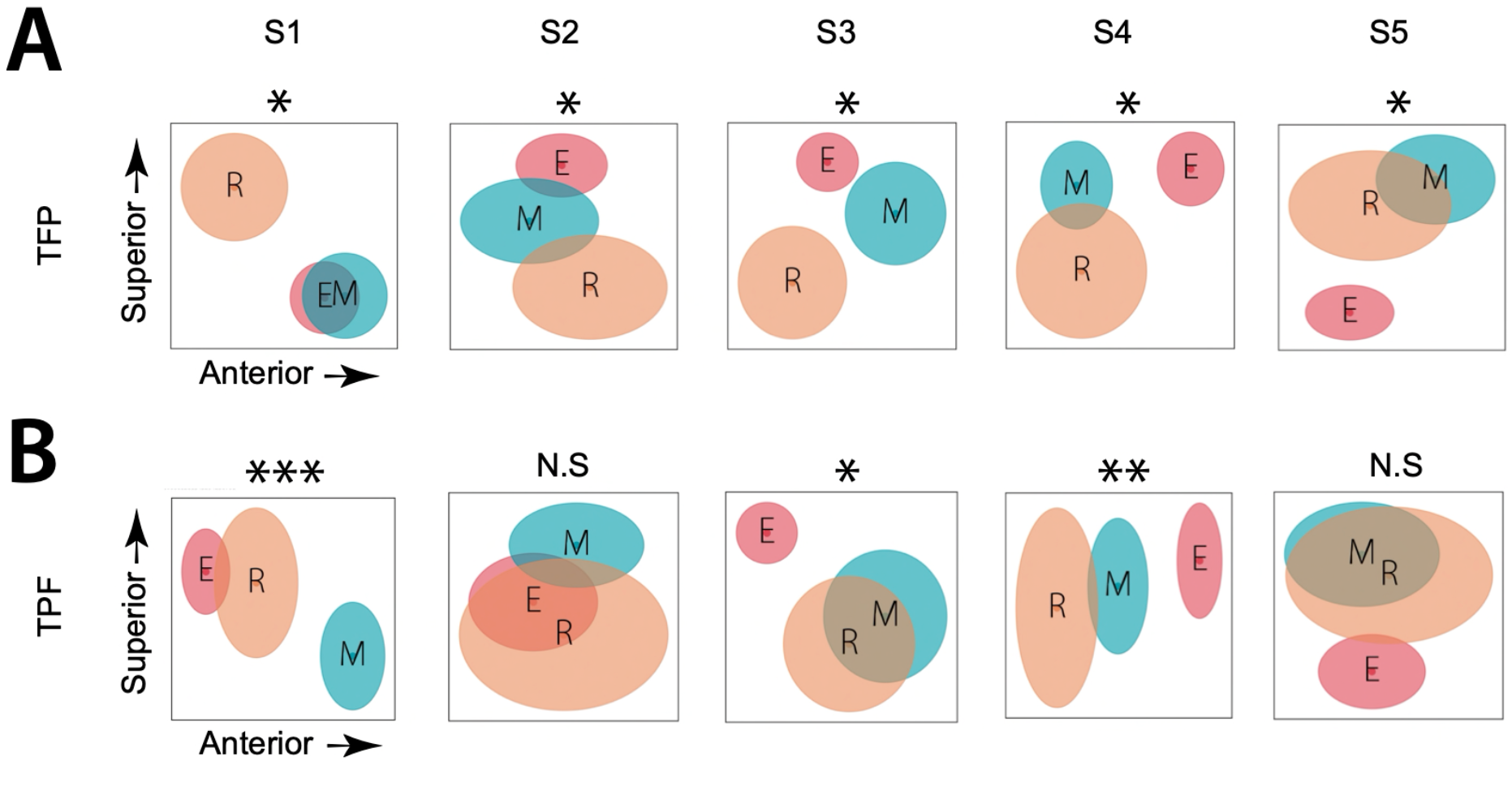
Group level results showing that the center of the TFP (A) and TPF (B) hidden spirals shift across the cortex to distinguish different behavioral states encoding (E), maintenance (M), and retrieval (R). **(A)** For encoding, N = 239, 220, 171, 188, 113 for S1-S5 respectively. For maintenance, N = 156, 110, 63, 149, 62 for S1-S5 respectively. For retrieval, N = 95, 95, 57, 54, 39 for S1-S5 respectively. **(B)** For encoding, N = 237, 109, 289, 212, 94 for S1-S5 respectively. For maintenance, N = 145, 81, 101, 140, 59 for S1-S5 respectively. For retrieval, N = 92, 37, 76, 68, 29 for S1-S5 respectively. For each ellipse (task period), the major axis (horizontal axis) denotes the standard-error-of-the-mean (SEM) for the Y-coordinate and the minor axis (vertical axis) denotes the SEM for the Z-coordinate, of the centers of the hidden spirals. (*** p < 0.001, ** p < 0.01, * p < 0.05, N.S. Not significant, MANOVA, FDR-corrected).

Thus, hidden spiral organization was not fixed to a particular anatomical location but dynamically reorganized with behavioral states. Importantly, this effect reflected changes in the spatial organization of the inferred dynamics, rather than simply changes in the strength or prevalence of traveling waves: the same cortical recording array expressed different underlying rotational configurations during different stages of memory processing. This state-dependent displacement suggests that hidden spirals may provide a flexible form of large-scale neural coordination whose spatial organization is coupled to the changing computational demands of memory. Although these findings do not establish a causal role for spiral relocation, they reveal a systematic relationship between latent cortical dynamics and behavioral state that is not apparent from the overt morphology of the recorded traveling waves alone.

## Discussion

Neural traveling waves exhibit diverse, complex spatial patterns, yet the dynamics that generate these patterns remain poorly understood. Here, we show that this apparent diversity can conceal a deeper layer of dynamical organization in human cortical activity. By modeling traveling waves as coupled phase oscillators while explicitly accounting for spatial heterogeneity in frequencies, we proposed FOCUS and identified hidden spiral dynamics within epochs whose recorded activity often showed planar, concentric, or complex spatial patterns. These findings suggest that the morphology of a traveling wave observed in neural recordings does not necessarily correspond directly to its underlying dynamical organization.

The FOCUS framework provides a mechanistic explanation for why overt spiral waves may appear relatively uncommon compared with plane and complex traveling waves. Coupled oscillator theory predicts that spatial heterogeneity in frequencies can generate plane and concentric waves and can strongly distort rotational dynamics. In a heterogeneous network, a latent spiral can therefore be expressed in the recorded activity as a spatially complex or apparently non-rotational pattern. Our observation that hidden spirals were preferentially associated with complex wave patterns, whereas stronger plane wave expression was associated with a lower probability of hidden spirals, is consistent with this theoretical framework. Rather than indicating that plane and spiral waves are independent phenomena, these relationships suggest that changes in spatial frequency structure can alter how an underlying dynamical organization is expressed at the level of observable wave morphology.

### Hidden spirals are distinct from overt rotational waves

An important question is whether the hidden spirals identified here are genuinely distinct from the rotational waves that can already be seen in neural recordings (Bhattacharya et al., 2022; Das et al., 2026; Prechtl et al., 1997), or whether our model simply recovers visible spirals in a transformed representation. Several observations argue strongly against the later interpretation. First, only approximately 10% of hidden spirals overlapped spatially with overt rotational waves identified directly from the recorded phase fields. Thus, most hidden spirals emerged from epochs that did not exhibit a corresponding visible spiral pattern. The hidden structures therefore cannot be explained simply as a re-detection of the rotational waves already apparent in the recordings. An overt spiral is a rotational organization that remains visible in the recorded phase field, whereas a hidden spiral is rotational organization whose spatial expression has been substantially distorted or obscured by heterogeneous local frequencies. This distinction is central to our interpretation: the model does not create a new wave morphology, but reveals dynamical organization that is not directly expressed in the observed morphology.

Several control analyses further support this interpretation (see **Methods** for details). First, randomizing the phases and then following the exact same computational analytical approach did not produce hidden spirals. Specifically, the same modeling and flattening procedure did not systematically produce spiral solutions when the relevant biological structure is removed or disrupted by using spatial autocorrelation-preserving surrogates on the phase data from the ECoG electrodes. Second, randomizing the original frequencies recovered from Eq. (2) did not produce spiral wave patterns. That is, when the frequency matrix was randomly selected from those recovered via Eq. (2) from phase data, with the selection probability weighted according to the task and subject-specific data distribution, the hidden spirals again vanished. And third, we used controls to ensure that our ECoG electrodes had sufficient spatial sampling to capture the hidden spirals from the biological structure of the neural data. Specifically, we generated simulated data using a bigger grid and then we subsampled phase data from the bigger grid with known ground truth hidden spiral and then followed the same modeling and flattening procedure on the subsampled data. This subsampled data preserved the hidden spirals that were present in the original bigger grid. Thus, the observed association between non-synchronous flattened solutions and hidden spirals cannot be attributed to an intrinsic bias of the chosen model. Rather, these controls support the interpretation that the hidden spiral geometry reflects genuine features of the underlying neural dynamics, rather than being a modeling artifact.

Importantly, identifying spatially or temporally structured components in neural recordings does not by itself reveal the dynamical mechanisms that generate traveling waves. Widely used data-driven approaches for dimensionality reduction, source separation, and latent-factor decomposition (Cunningham & Yu, 2014) are primarily designed to represent complex neural activity in terms of statistically distinct, low-dimensional, or otherwise structured components. Although these approaches can isolate prominent patterns of neural activity, they generally do not explicitly model the coupled phase dynamics among neural populations or incorporate local frequencies as mechanistic determinants of propagation. Consequently, a spiral wave that coexists with frequency-gradient-driven plane or concentric waves will not emerge as a distinct component; its activity will be distributed across multiple components, absorbed into dominant propagating modes, or represented as a mixture of spatial and temporal structure. Moreover, even when a decomposition identifies a component with rotational structure, it does not establish whether that structure arises from a dynamically organized phase singularity, local frequency heterogeneity, or the superposition of other propagating processes. Detecting hidden spiral dynamics therefore requires a framework that explicitly represents interactions among oscillatory neural populations, accounts for local frequency heterogeneity, and tests whether coherent rotational dynamics emerge after the effects of frequency-dependent propagation have been accounted for.

The distinction also explains why hidden spirals may be substantially more prevalent than suggested by direct observation. Spiral dynamics are particularly sensitive to spatial heterogeneity, and a spiral whose core lies outside the sampled cortical region can appear locally plane or spatially complex. Consequently, conventional analyses based solely on overt morphology may systematically underestimate the prevalence of rotational dynamics in cortical recordings.

### Hidden spirals provide a behaviorally structured dynamical organization

The hidden spirals were not spatially fixed. Their centers shifted across the cortical grid between encoding, maintenance, and retrieval, with TFP spirals showing significant state dependent displacement in all five subjects and TPF spirals in three subjects. This finding is important because it links the inferred latent dynamics to behavioral state rather than simply to the existence of traveling wave activity. The same cortical network therefore expressed different spatial configurations of rotational dynamics as memory processing progressed.

Such reorganization is consistent with the broader view that traveling waves provide a mechanism for dynamically coordinating distributed neuronal populations. By showing the existence of spiral traveling waves during human memory, our results reveal an additional new hierarchy of complex spatial patterns of traveling waves, complementing the traditionally known feedforward-feedback cortical hierarchy. Since traveling waves are known to be closely associated with spiking activity of neurons (Jacobs et al., 2007), their propagation putatively reflects packets of neuronal activity sequentially scanning distributed brain areas to transiently reorganize functional connectivity between them, to represent complex behaviors in human memory processing (Eichenbaum, 2000; Mesulam, 1990). In this context, it suggests that plane waves may be relevant for globally routing neural spiking activity, by directionally propagating the activity to different cortical areas in a feedforward-feedback manner. In contrast, because rotational waves revisit the same brain areas in multiple cycles, it suggests that these waves may be relevant for dynamically strengthening functional connectivity between large-scale neuronal assemblies for efficient memory processing and memory consolidation. The diverse propagation patterns of these cognition-related traveling waves are therefore consistent with a rich set of models, suggesting a role for spatially organized neural assemblies and oscillations in the computational processes underlying memory, cognition, and other behaviors (Freeman, 2003; Pinotsis et al., 2023; Pinotsis & Miller, 2023; Rubino et al., 2006). The state-dependent movement of hidden spiral centers therefore suggests that this dynamical organization can be flexibly reconfigured as the computational demands of memory change.

### Implications for the generation of cortical traveling waves

Together, our findings support a hierarchical view of traveling wave organization in which the spatial morphology observed in neural recordings represents only one level of the underlying dynamics. Local differences in frequency can shape the observable propagation pattern, while the coupled network can simultaneously support coherent rotational organization that is not overtly expressed. This framework provides a potential explanation for the striking diversity of cortical traveling waves without requiring every observed morphology to arise from a fundamentally different dynamical mechanism.

Given these factors, we modelled our spatiotemporally stable, complex spatial patterns using coupled phase oscillators, as traveling waves are oscillations whose phases move across the cortex to coordinate neural activity between different brain areas (Ermentrout & Kleinfeld, 2001). Coupled phase oscillator modelling has suggested that the changes in phases across a one- or two-dimensional domain can be due to spatial heterogeneities in frequencies. For example a linear spatial gradient in frequencies can lead to a plane traveling wave (Ermentrout & Kopell, 1984). Bullseyes or target waves can also be generated by frequency gradients using nearest-neighbor coupled oscillators (Ren & Ermentrout, 1998). Experimental data in humans has supported these modeling work by showing that frequency heterogeneities can produce plane waves (Zhang et al., 2018) and some complex patterns of traveling waves (Koller et al., 2024).

These heterogeneities often take the form of changes in the local frequency of the oscillators. However, certain spatial patterns of traveling waves such as rotating waves do not come from heterogeneities, rather they occur in uniform oscillatory networks (Ding & Ermentrout, 2022; Paullet & Ermentrout, 1994). These computational models have also suggested that, in the absence of any heterogeneities, synchrony and rotating waves are the only wave shapes that can occur as stable patterns in two-dimensional domains.

Even without the presence of any external sensory inputs, networks of homogeneously coupled phase oscillators can produce synchrony and spiral waves. However, if we propose that top-down and bottom-up signaling during the different task conditions of the working memory task do not alter the coupling (or, structural connectivity) between the oscillators, but rather alter the local frequency, then the oscillators are no longer homogenous. In the case of a fully synchronized homogeneous system, changes in the local frequencies will typically lead to patterns such as plane waves and target patterns. If the homogeneous network consists of a rotational or spiral wave, then heterogeneities due to external sensory inputs can dramatically distort the spiral wave. Thus, it may be that many complex spatiotemporal patterns that are recorded and observed in electrical recordings may actually be spiral waves that were distorted by extrinsic inputs perturbing the local oscillator frequencies. That is, rotational waves, rather than being somewhat rare, may, in fact, be much more prevalent than originally thought, but remain hidden. Consistent with this view, the hidden spirals that we detected were closely associated with larger traveling waves that we detected. Specifically, a presence of a plane wave in the stable epochs diminished the probability of detecting a hidden spiral, rather the hidden spirals were associated with complex spatial patterns of traveling waves, suggesting a new role of complex, heterogeneous spatial patterns in human memory processing and also suggesting that an intricate interplay between large, empirically observed traveling waves like plane waves and computationally detected hidden spirals.

Our results consequently expand the functional interpretation of cortical traveling waves during memory. Rather than being limited to the overt propagation patterns detected directly from recordings, the computational repertoire of traveling waves may include latent dynamical configurations that are selectively expressed, distorted, and spatially reorganized across behavioral states. Hidden spirals provide one example of this previously inaccessible layer of neural organization and suggest that the diversity of observed traveling waves may reflect, in part, the different spatial expressions of underlying dynamical processes rather than a collection of entirely distinct wave-generating mechanisms.

It is notable that the traveling waves that we detected often traveled across the sulci and gyri, and even across the Sylvian fissure, across distributed brain areas. Thus, the signals seemed to jump over sulci. For example, in **Figure 1C**, the spirals propagated through the Sylvian fissure. Recent work on computational modeling of neural field theory of brain waves (Pinotsis et al., 2023; Pinotsis & Miller, 2023) has shown that stable electric fields are capable of carrying information across sulci and gyri, thus providing a mechanism for ephaptic coupling of multiple, distributed brain areas. The traveling waves that we observed could putatively be a manifestation of these stable electric fields and their interactions with the sulci and gyri, which allows for ephaptic coupling between distributed brain areas (Ermentrout & Kleinfeld, 2001). An additional way that traveling waves could effectively propagate through sulci is by generation in subcortical areas, such as thalamus, as demonstrated by recent computational modeling work (Bhattacharya et al., 2021). Subcortical generation of traveling waves also hints at the possibility of three-dimensional traveling waves coordinating neural activity between the cortical and subcortical brain areas, rather than waves propagating only along the cortical surface. Denser sampling of depth electrodes in three-dimensions (Lubenov & Siapas, 2009) and three-dimensional computational modeling of traveling waves may help to better understand the full complexity of the generation and propagation of traveling waves across the widely varying shape of the cortex.

### Potential practical applications of hidden spirals

Hidden spirals could potentially be used as a biomarker for treatment of neurological and psychiatric disorders. Because spiral cores are phase singularities where the direction of oscillations wraps around a point, stimulating near these singularities (via TMS, tDCS, or intracranial electrodes) could allow precise control of large-scale wave propagation. In our working memory task, the hidden spirals shifted their centers to flexibly adapt to different behaviors such as encoding, maintenance, and retrieval, suggesting that the core or center of the hidden spirals is a dynamic, rather than static phenomenon, and helps the brain transition between different behavioral states.

Indeed, consistent with these empirical findings, computational modeling has suggested that spiral cores can be induced and modulated in uniform oscillatory systems by using delayed feedback and changing the feedback parameters (Hu et al., 2010). In vitro cardiac electrophysiology in rodents has suggested that using optogenetics, spiral cores can be precisely moved along pre-defined trajectories, thus demonstrating real-time spatiotemporal control over spiral wave dynamics in a biological system (Majumder et al., 2018). This offers a new way to redirect, stabilize, or suppress abnormal cortical activity. For example, closed-loop detection of hidden spirals could trigger targeted stimulation before symptoms manifest in memory disorders such as the Alzheimer’s disease. On the other hand, brain stimulation techniques can be tuned to entrain beneficial spirals, therefore supporting working memory, sequential processing, or error correction. For example, oscillatory tACS aligned with spiral phase progression could enhance temporal coding and long-range communication. Controlled spirals therefore could aid motor sequencing or language recovery by re-establishing timing scaffolds.

Furthermore, since spirals provide a distributed phase map across space, a brain-computer interface could read out these hidden dynamics to infer ongoing mental states with higher precision. This could enable spatial phase coding-based brain-computer interfacing, where information is extracted not just from power, but from spiral orientation and drift. Traditionally, brain stimulation targets locations or nodes for modulating neural activity. Hidden spirals provide us with new means by which we should target patterns (topological motifs), rather than nodes, therefore moving away from “node stimulation” to “wave geometry stimulation”, opening a new field of topological neuromodulation. In summary, the hidden spirals could act as a control point, where by detecting and modulating these hidden spirals, we could develop new interventions for psychiatric disorders and cognitive enhancement. Therefore, spirals could potentially represent a topological lever for stabilizing or reshaping global brain dynamics.

## Materials and Methods

### Human subjects

We examined direct brain recordings from 5 patients (N=5, 4 males, minimum age = 20, maximum age = 37, mean age = 29) with pharmaco-resistant epilepsy who underwent surgery for removal of their seizure onset zones. All patients consented to having their brain recordings used for research purposes and all research was approved by Institutional Review Boards. Recordings were collected at the Thomas Jefferson University Hospital, Philadelphia and the University of Pennsylvania Hospital Philadelphia. The direct recordings of these patients can be downloaded from https://memory.psych.upenn.edu/Data (Jacobs & Kahana, 2009).

### Electrophysiological recordings and preprocessing

Patients were implanted with different configuration of electrodes based on their clinical needs, which included both electrocorticographic (ECoG) surface grid and strips as well as depth electrodes. In this work, we only included patients who were implanted with 8×8 ECoG grids on the cortical surface. ECoG recordings were obtained using subdural grids (contacts placed 10 mm apart). Anatomical localization of electrode placement was accomplished by co-registering the postoperative computed CTs with the postoperative MRIs using FSL (FMRIB (Functional MRI of the Brain) Software Library), BET (Brain Extraction Tool), and FLIRT (FMRIB Linear Image Registration Tool) software packages. Preoperative MRIs were used when postoperative MRIs were not available. From these images, we identified the location of each recording contact on the CT images and computed the electrode location in standardized Talairach coordinates.

Original sampling rates of ECoG signals were 400 Hz, 512 Hz, and 1000 Hz. Therefore, ECoG signals were downsampled to 400 Hz, if the original sampling rate was higher, for all subsequent analysis. We used common average referencing (ECoG electrodes re-referenced to the average signal of all electrodes in the grid), similar to our previous studies on traveling waves (Das et al., 2022; Zhang et al., 2018). Line noise (60 Hz) and its harmonics were removed from the ECoG signals. For filtering, we used a fourth order two-way zero phase lag Butterworth filter throughout the analysis.

### Sternberg verbal working memory task

Patients performed multiple trials of a Sternberg verbal working memory task (Jacobs & Kahana, 2009; Sternberg, 1966). In each trial of the task, first the patients viewed a fixation cross and then they were sequentially presented with one to three English letters on the screen of a bedside laptop computer (*encoding*, **Figure 1A**). Each letter appeared on screen for 1 sec. Patients were instructed to closely attend to each stimulus presentation and to silently hold the identity of each item in memory (*maintenance*). After the presentation of each list, the subject viewed a probe item and was asked to respond by pressing a key to indicate whether the probe was present in the just-presented list or whether it was absent (*retrieval*). After the key press, the computer indicated whether the response was correct, and then a new list was presented. The 5 subjects completed 10 sessions in total. Mean accuracy across subjects was ∼ 91%. We analyzed the 1 sec long trials from the encoding period of each letter. For retrieval, we analyzed 1 sec long trials immediately after the subjects viewed the probe item. For maintenance, we analyzed 2 sec long trials in between the encoding and retrieval periods.

### Identification of oscillation frequency

Our first goal was to identify the oscillations that were present on multiple electrodes, and then (as described in subsequent sections), to map their spatial pattern of propagation as traveling waves. We focused our analysis on oscillations in the theta/alpha and beta bands. To identify the oscillations present on multiple channels, we adopted methods similar to our previous approach (Das et al., 2022; Das et al., 2025; Das et al., 2023). This subject-specific algorithm accounts for individual differences in electrode positions and oscillation frequencies across individuals. To identify traveling waves, a single oscillation frequency per electrode group is crucial since, by definition, a traveling wave involves a single frequency and whose phase progressively propagates through a contiguous region of cortex.

In this algorithm, to identify the frequency of interest for each electrode we first used Morlet wavelets to compute the power of the neural oscillations at 200 frequencies logarithmically spaced from 3 to 40 Hz. To identify narrowband oscillations, we fit a line to each patient’s mean power spectrum (mean across trials) in log–log coordinates using robust linear regression (Das et al., 2022). We then subtracted the actual power spectrum from the regression line. This normalized power spectrum removes the 1/f background signal and emphasizes narrowband oscillations as positive deflections. We identified narrowband peaks in the normalized power spectrum as any local maximum greater than one standard deviation above the mean. The mean of the peak frequencies of the electrodes of a ECoG grid was defined as the grid’s cluster frequency (CF). Overall, ∼ 80% of ECoG electrodes had a narrowband oscillation in one of the theta/alpha (< 12 Hz) or beta (> 12 Hz) frequency bands, indicating that an overwhelming number of ECoG electrodes show oscillatory activity. Since all electrodes in an oscillation cluster will not have identical oscillation frequency, we also allowed for electrodes to show oscillations at nearby but nonidentical frequencies (within 5 Hz of cluster frequency). Our traveling waves analysis below was carried out based on the individual frequency of each electrode to account for the variation of frequencies across electrodes.

### Identification of traveling waves

Our next objective was to identify and characterize traveling waves. A traveling wave can be described as an oscillation that moves progressively across a region of cortex. Quantitatively, a traveling (phase) wave can be measured as a set of simultaneously recorded neuronal oscillations at very similar frequencies whose instantaneous phases vary systematically with the locations of the recording electrodes. To identify these traveling waves we used a localized circular-linear regression approach, assuming that the relative phases of the oscillation clusters exhibit a linear relationship with electrode locations *locally* (Das et al., 2022). This locally circular-linear fitting of phase-location can detect complex patterns of traveling waves in an oscillation cluster in addition to plane traveling waves.

To identify traveling waves from the phases of each oscillation cluster, we first measured the instantaneous phases of the signals from each electrode of a given cluster by applying a 4^th^-order Butterworth filter at the cluster’s oscillation frequency (bandwidth [f_p ×_.85, f_p /_ .85] where f_p i_s the peak frequency). We used the Hilbert transform on each electrode’s filtered signal to extract the instantaneous phase.

Next, we used circular statistics to measure the spatial propagation of phase between neighboring electrodes in each cluster to identify traveling waves at each time point (Fisher, 1993). To simplify visualizing and interpreting the data, we first projected the 3-D Talairach coordinates for each cluster to a 2-D plane using principal component analysis (PCA).

To identify traveling waves, we used a series of two-dimensional (2-D) localized circular–linear regression that were fit to model the direction of wave propagation in a local *subcluster* in the 25-mm neighborhood of each electrode. The local regression determines the direction of local wave propagation in the subcluster surrounding each electrode, by measuring whether the local phase pattern varies linearly with the electrodes’ coordinates in 2-D. Thus, this regression fits a single direction of wave propagation locally around each electrode at each moment, and these measures are then subsequently combined across subclusters to identify larger-scale spatial patterns of propagation.

In this regression, for each nearby electrode, we first identified the neighboring electrodes that were located nearby, constituting a sub-cluster of the given cluster. Here, let *x_i_* and *y_i_* represent the 2-D coordinates and *θ_i_* the instantaneous phase of the *i*th electrode in a sub-cluster. We used a 2-D circular-linear model

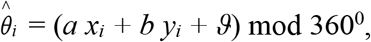

where 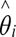 is the predicted phase, *a* and *b* are the phase slopes corresponding to the rate of phase change (or spatial frequencies) in each dimension, and *ϑ* is the phase offset. We converted this model to polar coordinates to simplify fitting. We define *α* = atan2(*b, a*) which denotes the angle of wave propagation and 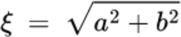 which denotes the spatial frequency. Circular–linear models do not have an analytical solution and must be fitted iteratively (Fisher, 1993). We fitted *α* and *α* to the distribution of oscillation phases at each time point by conducting a grid search over *α* ∈[0^0^, 360^0^] and ζ ∈ [0,18]. Note that ζ = 18 corresponds to the spatial Nyquist frequency of 18^0^/mm corresponding to the spacing between neighboring electrodes of 10 mm.

In order to keep the computational complexity tractable, we used a multi-resolution grid search. We first carried out a grid search in increments of 5^0^ and 1^0^/mm for *α* and ζ, respectively. The model parameters (*a= ξ*cos(*α*) and *b= ξ*sin(*α*)) for each time point are fitted to most closely match the phase observed at each electrode in the sub-cluster. After having relatively coarse estimates of *α* and ζ, we then carried out another grid search in increments of 0.05^0^ and 0.05^0^/mm around a ± 2.5^0^ and ± 0.5^0^/mm neighborhood of the coarse estimates of *α* and ζ, respectively, to have refined estimates of *α* and ζ. We computed the goodness of fit as the mean vector length *r̅* of the residuals between the predicted (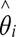) and actual (*θ_i_*_)_ phases (Fisher, 1993),

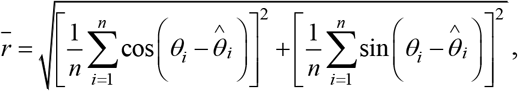

where *n* is the number of electrodes in the sub-cluster. The selected values of *α* and *ξ* are chosen to maximize *r̅*. This procedure is repeated for each sub-cluster of a given oscillation cluster. To measure the statistical reliability of each fitted traveling wave, we examined the phase variance that was explained by the best fitting model. To do this, we computed the circular correlation *ρ_cc_* between the predicted (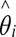) and actual (*θ_i_*_)_ phases at each electrode:

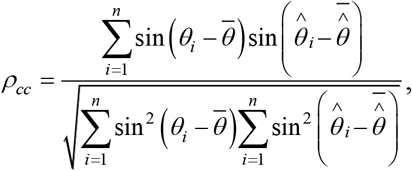

where bar denotes averaging across electrodes. We refer to 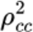 as the wave strength (Das et al., 2022) as it quantifies the strength of the traveling wave (note that 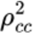 has been referred to as the phase gradient directionality (PGD) in some prior studies (Rubino et al., 2006; Zhang et al., 2018)). Other features of traveling waves such as the wavelength (2π/spatial frequency) and the speed (wavelength × frequency) can be readily derived from the parameters of the above 2-D model. Consistent with earlier work (Das et al., 2022; Halgren et al., 2019; Zhang et al., 2018), the mean propagation speed of the traveling waves in our dataset was ∼ 0.7 m/s.

Note that traveling waves in some prior studies were detected and analyzed by calculating the spatial gradient of the phases of the recordings from ECoG electrodes (Halgren et al., 2019), however, phase gradients can only be calculated in two directions (forward and backward), so only a subset of neighboring electrodes of a given electrode are included in these analyses of spatial gradient. Since our approach directly includes all possible neighboring electrodes (termed as a sub-cluster in our analysis) in the circular-linear regression model, thus calculating phase gradients in all possible directions, it results in a more efficient estimate of the traveling waves parameters.

We note that because a few of the ECoG electrodes did not have a narrowband oscillation on certain trials, to allow us to estimate the overall shape of the traveling wave across the cluster, we estimated the traveling waves for those electrodes by an extrapolation procedure where, for the given electrode, we substituted the mean of the traveling waves of all electrodes within a 25 mm radius (subcluster) of the electrode under consideration. This extrapolation step was necessary for classification of each oscillation cluster as one of the wave categories using curl and divergence analysis (rotational or expanding/contracting or complex, see section below on ***Identification of plane, rotational, and concentric traveling waves***). This is reasonable since as mentioned above, an overwhelming (∼ 80%) number of ECoG electrodes showed oscillatory activity. Nevertheless, we reran our complete series of analyses without the extrapolation step, and found that the same modes and clusters were still statistically significant, indicating that this step had very minimal effect on our results.

### Identification of stable epochs

Since a traveling wave is composed of phase patterns that vary relatively smoothly across space and time, we sought to characterize the spatiotemporal stability of the traveling waves that we detected in our localized circular-linear regression approach above. We defined *stability* as the negative of the mean (across all electrodes in an ECoG grid) of the absolute values of the difference between the direction and strength of traveling waves observed at consecutive time-points, defined as

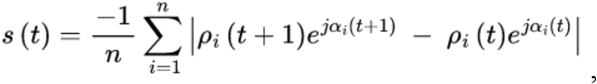

where, *ρ_i_* denotes the strength of the traveling wave for the *i*^th^ electrode, α*_i_ n* denotes the direction of the traveling wave for the *i*^th^ electrode, *n* denotes the number of electrodes in an ECoG grid, *s* denotes stability, *t* denotes time, and *j* denotes square root of minus one. We repeated this procedure for each pair of consecutive time-points and for each trial of the working memory task. Stability for each trial was z-scored. We identified stable epochs as those for which all stability values were above the mean of the stability values for a given trial.

Previous research in rabbit field potential recordings (Freeman & Rogers, 2002; Freeman & Schneider, 1982) have found spatial patterns of theta traveling waves last ∼80-100 msec in duration. Moreover, large-scale, whole-brain computational modeling in humans using neural field theory have shown that spatiotemporally stable traveling waves last ∼50-60 msec in duration (Roberts et al., 2019). Therefore, we focused our analyses on the stable epochs that were more than 50 msec long for further analysis; as there was evidence that shorter wave patterns may be less stable (Roberts et al., 2019). In our dataset, mean length of the stable periods was ∼ 143 msec across subjects.

### Identification of modes using complex independent component analysis (CICA)

We next used a large-scale dimensionality reduction framework to measure the spatial patterns of wave propagation across each cluster. This framework identified the different types of spatial wave patterns that formed traveling waves at each cluster and modeled how they summed to determine the signal on each epoch. We thus applied complex ICA (CICA) to the data on each epoch. Within each stable epoch identified as described above, the direction and strength of the traveling waves remain almost consistent across time-points. Thus, we averaged the direction and strength of the traveling waves within each stable epoch to find one spatial wave pattern associated with each stable epoch, which was input to the CICA algorithm.

Next, to analyze how the spatial patterns of waves changed over time, we concatenated the wave patterns (i.e., direction and strength) across all stable epochs into a single matrix and then passed this matrix as input to the complex version of the independent component analysis (CICA) (Fu et al., 2015; Li & Adalı, 2010). We used this complex version of the ICA, as compared to real ICA, to incorporate the 2-D directions of the traveling waves, weighted by the strength (ρ^cos^ (α) and (*α*), defined for each electrode in the ECoG grid. We then extracted the independent activation functions (or, weights) (each activation function corresponds to one of the stable epochs) and the corresponding modes (**Figure 2**) as the output from the CICA (Fu et al., 2015; Li & Adalı, 2010). Multiplication of each of these modes with the mean of the weights across all stable epochs corresponding to that specific mode results in a unique wave pattern (termed as *mean mode*) associated with that mode. At the individual epoch level, a higher CICA weight for that epoch corresponding to a specific mode indicates higher representation of that wave pattern in that specific epoch and a lower CICA weight for an epoch corresponding to a specific mode indicates lower representation of that wave pattern in that specific epoch (**Figure 2**). Moreover, the higher the variance explained by a given mode, the higher will be its representation across the trials. Generally, we observed more complex wave patterns in higher order modes (modes beyond 3), which suggests that complex wave patterns explain relatively less variance in the data compared to plane and spiral wave types. On average the first three modes explained ∼27%, ∼17%, and ∼13% variance respectively, and the first eight modes combined explained >80% variance. In this way, we can extract the CICA weights for each of the encoding, maintenance, and retrieval periods, and compare them statistically, see ***Statistical analysis*** section below.

### Identification of planar, rotational, and concentric traveling waves

After extracting the mean modes from the CICA procedure above, we next sought to classify each of the mean modes into one of the following categories “planar”, “rotational”, “concentric” (“expanding” or “contracting”), or “complex” (**Figure 2**) (Das et al., 2026), to identify global patterns of traveling waves in particular shapes that might be meaningful for cortical processing.

For classifying each of the modes into a specific wave pattern, we carried out robust statistical tests. We constructed appropriate surrogate distributions by shuffling the coordinates of the electrodes which destroys the specific structure associated with a given wave pattern.

For classifying the specific shape of each wave, we created a series of scores, which identified plane waves, and rotational/concentric waves. Clusters that did not meet our criterion for planar/rotational/concentric waves were labeled as ‘complex’. For detecting plane traveling waves, we calculated the mean wave direction (weighted by the strength of the wave) of all electrodes in an oscillation cluster. We then constructed surrogate distributions by shuffling the coordinates of the electrodes and recalculating the mean wave direction for each shuffling, therefore constructing a null distribution against which the empirical data can be compared.

From the surrogate distribution, the threshold that corresponded to significance at *p* = 0.001 was ∼ 0.6. Thus, we declared a wave as a plane wave for which the mean wave direction was higher than 0.6. Visually, this procedure of threshold selection identified individual mean modes that showed clear plane waves (**Figures 2**, **9**) and small changes in this threshold did not substantially change the results.

Similarly, we classified the mean modes that represented rotational and concentric traveling waves by measuring the shape of their wave propagation using the *curl* and *divergence* metrics respectively (Das et al., 2026). Curl measures rotational patterns (for example, clockwise or counter-clockwise rotation) in wave dynamics and divergence measures expanding/contracting patterns (for example, sources or sinks) in wave dynamics (Das et al., 2026). We first calculated the curl and divergence metrics for each electrode and then calculated the mean curl and divergence across all electrodes in an oscillation cluster.

The curl of a traveling wave at a particular electrode can be calculated as follows. For a 2D traveling wave 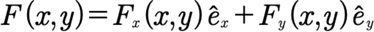 at electrode locations (*x, y*), where *F_y_ (x, y)* and *F_y_ (x, y)* denote the traveling wave components in *x* and *y* directions respectively and *ê_x_* ê_x_ and *ê_y_* denote the unit vectors in the *x* and *y* directions respectively, curl can be defined as

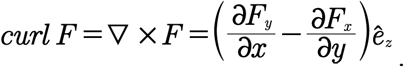

Similarly, the divergence of a traveling wave at a particular electrode can be calculated as follows

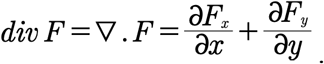

We used the same approach to assess significance of these labels as above, constructing appropriate surrogate distributions by shuffling the coordinates of the electrodes and then recalculating the curl and divergence. For *p* = 0.001, we identified curl and divergence thresholds of ∼ 0.75 and ∼ 0.4 for the rotational and concentric waves, respectively. We thus declared a wave as a rotational or concentric wave for which the curl or divergence value was higher than 0.75 or 0.4 respectively. Visually, this procedure of threshold selection functioned as we expected, as the waves detected with these thresholds showed clear rotational and concentric dynamics visually (**Figure 2**) and small changes in this threshold did not substantially change the results reported here.

Modes that were not classified as one of the planar, rotational, or concentric, were designated as complex waves. Complex waves were also stable at the individual epoch level and also across epochs, as these waves were able to distinguish behavior (**Figure 3**). Even though there was no global pattern associated with these complex traveling waves, many of these complex waves showed interesting local patterns, where a subset of electrodes in an ECoG grid showed planar, rotational, or concentric waves (**Figures 2**, **3, 9**).

### Coupled phase oscillator modeling

Since the complex spatial patterns of traveling waves that we detected were from phases extracted from oscillations present in the ECoG electrodes, we used coupled phase oscillators as a model for spatiotemporally stable traveling waves (Schwemmer & Lewis, 2011). Indeed, previous research has shown that two-dimensional arrays of nearest neighbor coupled phase oscillators can produce rotating and also complex spatial patterns of traveling waves as stable patterns, including in experimental data (Ding & Ermentrout, 2022; Koller et al., 2024; Paullet & Ermentrout, 1994). We applied coupled phase oscillator modeling on the spatiotemporally stable epochs that were present in the individual trials of the working memory task and across task conditions such as the encoding, maintenance, and retrieval.

A natural model for such data is the spatial Kuramoto phase model on a two-dimensional grid (Kuramoto, 1984):

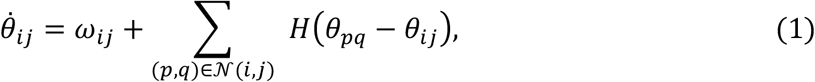

where *θ_i_*_j_ ∈ T = [0,2*π*) is the phase at electrode (*i*, *j*), *ω_i_*_j_ is the natural frequency, coupling is with the nearest neighbors N(*i*, *j*), and *H*(*φ*) is a 2*π* periodic interaction function. We have made several simplifying assumptions to minimize the number of free parameters. Specifically: (1) interactions are nearest neighbor, (2) the only difference between the oscillators is their natural frequency, *ω_i_*_j_. In addition to choosing the frequencies, we must choose an appropriate form for *H*(*φ*). We discuss this below.

### Recovering frequencies from phase data

Since during the stable epochs, the relative phases between the electrodes remain almost constant across time, we assume that the solutions to Eq. (1) are phase-locked, that is, they all have the same ensemble frequency, Ω. That is, *θ_i_*_j_(*t*) = Ω*t* + *φ_i_*_j_ with offsets *φ_i_*_j_ constant and determined from the data frame. In order to eliminate the dependence on Ω, we replace *θ_i_*_j_ by *θ_i_*_j_ − *θ*_11_, which are the phases relative to the oscillator at (1,1). Any electrode can be chosen as the reference electrode, without changing the phase-locked solution. The choice of (1,1) is for convenience. Thus, to get *φ_i_*_j_ from the data frame, we take *φ_i_*_j_ = *θ_i_*_j_ − *θ*_11_. Substituting into Eq. (1) yields:

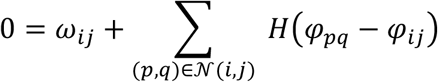

Hence the frequency at each node can be extracted from the phase *φ* via

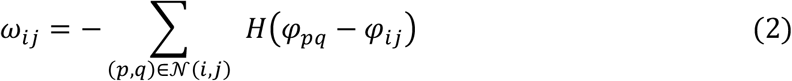

It is worth noting that *ω_i_*_j_ depends on the choice of the coupling function *H*(*φ*). Finally, if all the *ω_i_*_j_ are identical, then *synchrony*, *φ_i_*_j_ = 0 is always a solution to Eq. (2).

### Dynamical stability

In order for a solution for a differential equation such as Eq. (1) to make physical sense, it has to be a stable equilibrium (Strogatz, 2024). To distinguish from the notion of stability for the experimental data, where the same pattern of phases persisted over several frames, we refer to this as dynamical stability. To study dynamical stability of a phase-locked solution 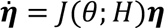, we linearize (1) about *θ* to obtain ***η̇*** = *J*(*θ*; *H*)***η***, where *J* is the Jacobian matrix of partial derivatives of the right-hand side with respect to phase-differences evaluated at the locked phases, *φ_i_*_j_.

Stability requires that *J* have no eigenvalues with positive real part (Strogatz, 2024). The entries in *J* consist of terms proportional to *H*^′^(Δ*φ*) where *H*^′^(*φ*) is the derivative of *H*(*φ*) and Δ*φ* are the phase-differences between adjacent electrodes. We now make our first constraint on *H*(*φ*).

From the previous section, if all the frequencies are identical, then synchrony is a phase-locked solution. We assume that if synchrony is a locked solution, then it is dynamically stable. This means that it must be the case that *H*^′^(0) > 0 (Ermentrout, 1992). For general phase-locked solutions, the more positive the numbers *H*^′^(Δ*φ*) then the more negative are the eigenvalues of *J* (Ermentrout, 1992). Since *H*^′^(0) > 0, this means that if the phase-differences between adjacent electrodes are small enough, then *H*^′^(Δ*φ*) > 0 and we will get dynamic stability. On the other hand, if the phase-differences are larger than roughly *π*/2, it is likely that *H*^′^(*φ*) < 0 and there is a bigger chance for dynamical instability (Ermentrout, 1992). Because the ECoG electrodes are relatively widely spaced, the phase differences can be quite large, which increases the chance that the patterns will be dynamically unstable. Hence, we need to embed the experimental array of phases in a larger network to create smaller phase differences.

### Interpolation to reduce phase gaps

To reduce phase gaps between electrodes in an 8 × 8 ECoG array, we introduce additional nodes (shadow oscillators) at intermediate lattice sites, first forming a 15 × 15 grid and then subsequently interpolating the 15 × 15 grid to a 29 × 29 grid (**Figure 4**). Since the distance between the adjacent ECoG electrodes is 10 mm and the diameter of each of the ECoG electrode is 2.4 mm, a 29 × 29 interpolation is essentially equivalent to physically packing three additional electrodes in between each of the adjacent ECoG electrodes (**Supplementary Figure 2**).

We then solve the following differential equation until it reaches a steady-state (constant phase-differences):

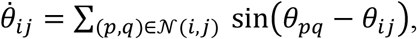

for all the new node phases that we have introduced, while keeping the *θ_i_*_j_ from the data values frozen at their original values. This produces a smoothly interpolated phase field with reduced nearest-neighbor gaps, whose equilibrium we use for analysis. We emphasize that the phases at the original electrodes are identical to those taken from the data.

### Estimation of coupling function

Since the traveling waves are oscillatory, we used an oscillatory coupling function. We determined the coefficients of this coupling function using a constrained, gradient-based method. Each coefficient in the coupling function denotes the relative contribution of the terms for a particular stable epoch. Once the coupling function is determined, the frequency values can then be recovered by solving the Kuramoto oscillator model, using the phases of the oscillators corresponding to the stable epoch.

In this paper, we use *H*(*φ*) of the following form

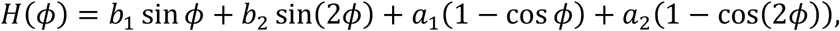

where coefficients *a*_j_, *b*_j_ can be tuned separately for each stable epoch to meet dynamical stability requirements. We also assume that *b*_1_ > 0 so that synchrony is dynamically stable when it is a solution (see above discussion), therefore, without loss of generality, we set *b*_1_ = 1. For a given phase network *φ*, we choose the (*b*_2_, *a*_1_, *a*_2_) triplet to increase the linear stability margin by minimizing the real parts of the eigenvalues of the Jacobian matrix, *J* :

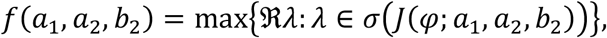

where *σ* is the spectrum of the matrix. We want to make *f* as negative as possible to increase the linear stability margin. After setting initial guess and parameter bounds, we run a constrained, gradient-based method (sequential quadratic programming or SQP) to sequentially update the parameters that reduce *f* while staying inside the bounds until the iteration is stopped by test triggers. Once *H* is determined, *ω_i_*_j_ values can then be estimated from the phase values using Eq. (2).

### Flattening method to reveal underlying patterns

If all the frequencies of the oscillators are the same, then, as we have noted, synchrony is a dynamically stable solution. However, it is not the only stable solution to a network of identical nearest-neighbor coupled phase oscillators. Among the other classes of stable solutions are spiral or rotating waves (Paullet & Ermentrout, 1994). Both synchrony and rotating wave patterns become very distorted and appear as heterogeneous, complex spatial wave patterns when there are frequency heterogeneities; therefore, to reveal the underlying patterns without the heterogeneities, we have devised a novel algorithm FOCUS (**F**requency-flattened **O**scillators in **C**ortex **U**nmask **S**pirals), to flatten the frequencies (**Figure 6**).

We pick a number of steps, *N_f_*, typically *N_f_* = 80, and let ℎ = 1/*N_f_*. We then solve the following differential equation until it reaches a steady-state phase-locked solution:

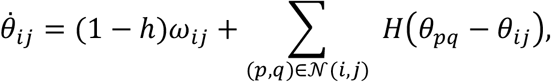

with initial condition, *θ_i_*_j_(0) = *φ_i_*_j_, the phases from the interpolated data. And *ω_i_*_j_ are the frequencies recovered with Eq. (2) from the phase data. We call the steady-state of this equation, 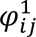. We then iteratively solve:

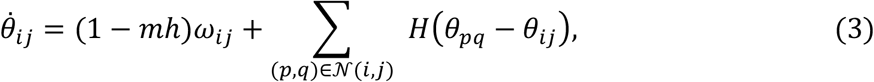

for *m* = 2,3, …, *N_f_*, with 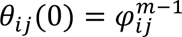 until a steady state is reached, and call this steady state, 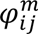. Once *m* = *N_f_*, the frequencies of all the oscillators are identically 0 since 1 − *N_f_*ℎ = 0. This procedure is an approximate method to continue along branches of solutions as a parameter (here, 1 − *m*ℎ) changes (Krauskopf et al., 2007). The steady-state solution 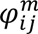 is the underlying *hidden* phase pattern. Intriguingly, all the non-synchrony solutions in our experimental data were spirals, suggesting that spirals form a *core* or *basis* wave pattern.

To rule out the possibility that the hidden spirals could have been a result of the rotational wave patterns present in the original stable epochs, we calculated the amount of overlap between the rotational or spiral waves at the individual stable epochs with the hidden spirals estimated from those same stable epochs. We detected rotational waves in the individual stable epochs using our curl measure (see the section ***Identification of plane, rotational, and concentric traveling waves***). This analysis revealed only ∼ 10% overlap between the rotational waves and the hidden spirals across task conditions and subjects, suggesting that FOCUS reveals dynamical organization that is not directly expressed in the observed morphology.

There are a couple of technical points that are worth pointing out. First, because we want to keep tracking the stable branch of solutions during the flattening process, we require the solution of each forward ODE to be stable to continue. If during the process the solution of one step is dynamically unstable, then the process has to be terminated, and the results remain inconclusive. Second, by the nature of the multi-stability of solutions, it is possible to jump to a different set of phase-locked patterns (**Figure 5**). To ensure our results fall on the same solution branch that our initial condition started on, we simply reverse the process. That is, we start from the pattern obtained from the flattening process and gradually increase the frequency heterogeneity by solving the same ODE sequence in the opposite order. We then compare this converged phase pattern, where the frequency is at its original level, with the interpolated phase pattern we started with. If those phase networks match, we conclude that the results are robust. Overall, we found that ∼ 23% of all stable epochs in our experimental data were dynamically stable and robust, with ∼ 10% hidden spirals and ∼ 13% synchrony. These numbers were statistically robust, see ***Statistical analysis*** section for more details.

### Continuation

To further ensure that the results remain on the same solution branch without jumping to a different set of phase-locked patterns, we further applied a pseudo-arclength continuation procedure to the same family of flattened-frequency equilibria (Govaerts, 2000). This numerical method allows us to trace solution branches of nonlinear equations, including through folds or turning points where standard continuation may fail. Using the interpolated phase pattern θ, we recovered the frequency field ω with Eq. (2) from the equilibrium condition and considered the branch defined by:

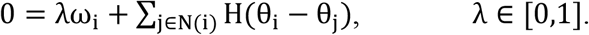

Here *λ* = 1 gives the original full frequency heterogeneous solution and *λ* = 0 gives the fully flattened underlying pattern. To seed the continuation branch, we solved for a few nearby equilibria at slightly smaller *λ* values using Newton iterations. And then we advanced it through branch with predictor-corrector pseudo-arclength method with adaptive step size (Govaerts, 2000). We reject iteration candidates if one of the following scenarios occur: (i), Newton corrector failed; (ii), branch exhibited an abrupt phase jump; or (iii), iteration lost dynamical stability.

### Identifying the chirality, core, and twist of the hidden spirals

In a phase network, a spiral core or center can be identified as a 2 × 2 block of oscillators where the phase advances by approximately an even 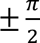 step along each edge, such that the four edges together complete an approximate ±2*π* winding around the block. The chirality or orientation of the spiral is then determined by the direction of this even quarter-turn winding: clockwise for 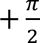 steps and counter-clockwise for 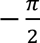 steps.

To detect such structures, we slide a 2 × 2 window over the phase matrix

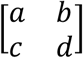

and compute the 2*π*-wrapped phase differences along the loop

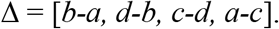

For each difference, we evaluate its chord distance on the unit circle to the expected quarter-turn, either 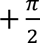 or 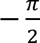, using the order parameters

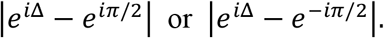

The root-mean-square of these distances provides a local mismatch score, and blocks whose score falls below a predefined threshold are classified as spiral cores. The sign of the winding then specifies the chirality or orientation, allowing us to detect both the position and the chirality of spiral centers across the lattice (**Figure 7**).

Converting the orientation to anatomically relevant terms, we refer to clockwise/counter-clockwise spirals as temporal◊frontal◊parietal (TFP) or temporal◊parietal◊frontal (TPF) respectively. Additionally, we also quantified the degree of the twist (T) of the hidden spirals (**Figure 8B**). To quantify the twist of a spiral, we used the adapted metric 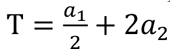, where *a*_1_and *a*_2_ are the coefficients of cos *φ* and cos 2*φ* in the coupling function respectively. This metric is obtained by expanding the two cosine terms to second order in *φ* and then combining their quadratic contributions. We also verified through simulations that this metric is aligned with the observed degree of spiral twist. The sign (positive/negative) of the twist metric T indicates whether it’s an inward (spiral-in) or outward (spiral-out) hidden spiral respectively (Ermentrout, 1995) (**Figure 8A**).

### Effect of interpolation on the detection of hidden spirals

#### Control analysis 1

In order to rule out the possibility that our sine-based interpolation procedure spuriously generates spiral patterns, we conducted control analyses based on forward-integration. We initialized an 8 × 8 phase lattice in a synchronized state. For each trial, a frequency matrix was randomly selected from those recovered via Eq. (2) from phase data, with the selection probability weighted according to the task and subject-specific data distribution. Using the same coupling function *H*(*φ*) like previously, we then solve the following forward ODEs:

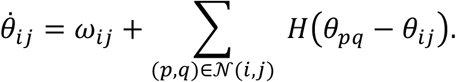

Then we interpolate the 8 × 8 solution to a 29 × 29 phase network and subsequently remove frequency heterogeneity from the simulated phase patterns using a flattening procedure before counting spiral cores. Across 200 trials, we observed spiral formation in only 1 case (0.5%), indicating that spurious spirals solely due to the interpolation step are extremely unlikely under these conditions.

#### Control analysis 2

To evaluate whether the flattening procedure preserves the underlying spatial phase pattern after coarse sampling and interpolation, we performed a validation test on randomly generated phase equilibria. We first generated 29×29 phase patterns from random initial conditions in a system equipped with random data-like scaled frequencies, symmetric 4-nearest-neighbor coupling, and a pure sine coupling function. The system was integrated until a stable equilibrium phase P was reached. For each equilibrium pattern P, we then applied the same flattening algorithm used in the phase data analysis. The flattened result from the original 29×29 pattern was labeled Group A.

We next extracted an evenly spaced 8×8 sub grid from the same 29×29 equilibrium, interpolated it back to 29×29 using the same sine interpolation. We then flattened the interpolated pattern again. The flattened result was labeled Group B.

We then compared the final Group A and Group B patterns, assessing agreement by comparing for both the spiral and sync classifiers and also core locations. We repeated this procedure 50 times. This analysis revealed that only 1 out of the 50 trials has disagreement between the 2 groups, corresponding to 98% accuracy. This analysis shows that hidden spirals can still be recovered from spatially subsampled data.

### Delay models

Delay models have been widely employed in the study of neuronal networks, as they provide mechanistic explanations for a variety of spatiotemporal phase pattern formations. This is particularly relevant in the case of unmyelinated axons, where signal transmission is relatively slow. For weakly coupled oscillators with delay, we have:

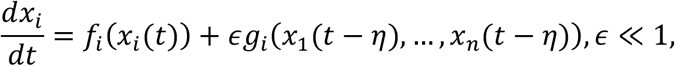

where *f_i_* are the frequencies, *g_i_* are the interactions received, and *η* is the delay. This system can then be reduced to a simpler phase description (Ermentrout, 1994):

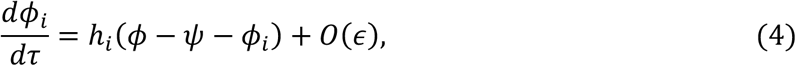

where ℎ*_i_* are the phase interaction functions, *φ* = (*φ*_1_, …, *φ_n_*)^T^ ∈ T*^n^*, T*^n^* is the *n*-torus. In such reduced phase models, delay terms manifest as the phase shift *ψ* within the argument of the interaction function. However, as shown in (Izhikevich, 1998), an explicit time delay emerges in the phase description only when the transmission delay is sufficiently large.

In the context of the present study, all our interactions are nearest neighbor, so that the delay between connected oscillators is identical for all interactions. As these delays are short relative to the period of the oscillations, they become phase-shifts of the interaction function (see Eq. (4)). Phase-shifts just change the coefficients of the sine and cosine terms and our choice of interaction functions already contain both sine and cosine terms. Hence, adding delays would be redundant as the phase-shifted interaction functions are already included in our model.

### Alternative dynamical models

While we have used the Kuramoto phase model for modeling our stable periods of traveling waves, we also have considered other dynamical models and mechanisms that could have potentially been used for our memory data analysis, however these were excluded for reasons outlined below.

#### Wilson-Cowan (WC) model and Jansen-Rit (JR) model

Neural field models such as the **Wilson-Cowan (WC) model** (Ermentrout & Cowan, 1979; Wilson & Cowan, 1972) with excitatory and inhibitory populations or the three-population **Jansen-Rit (JR) model** (Jansen & Rit, 1995) are often used to generate oscillatory behavior over a wide range of frequencies and varying the drive can induce frequency differences. Indeed, the JR model with a spatially-dependent drive has been used to produce traveling plane waves in the hippocampus (Behler et al., 2026). By coupling networks of these oscillators in a spatial grid, it is possible to produce a variety of wave patterns including sources, sinks, plane waves, and even rotating waves. However, in order to explain the data on our electrode arrays, we have the “inverse problem”: determine the drives and the connectivity between modules to produce a *particular* pattern of phases. While it is reasonably straightforward to compute the dependence of the frequency in a local module on the drive, there is no clear way to translate a combination of frequency differences and coupling between the modules to a give phase (or timing) difference. For example, in the WC equations, there are four different ways to couple modules: E◊E, E◊I, I◊E, and I◊I, all of which lead to different phase lags between the ongoing oscillations. In a single mutually coupled pair, then, it would be possible to tune coupling to attain a given lag.

However, in a network, one has to vary coupling between N×N modules, where N is the number of electrodes in an ECoG grid; changing the coupling between one pair of modules also changes the phase lags between that module and all its other neighbors. This *synergistic* interaction makes it impossible to tune coupling to match the whole electrode pattern.

#### Anisotropic coupling

Our rationale to use frequency gradients to generate plane waves was because of its straightforward interpretation. Frequency gradients arise naturally as the stimulus to the network is varied. Therefore, the fact that local frequencies can change based on external perturbation is very much compatible with the changes in top-down or bottom-up drive to whichever brain regions are generating the oscillations. That is, one can keep exactly the same network connectivity but by altering the drive to the neural units, one can manipulate the local frequency that can then drive the changes in the parameters of the phase model. However, frequency gradients are not the only mechanism to generate plane waves, there are other possible mechanisms to generate plane waves and spiral waves. However, in our dataset we observed a diverse range of complex spatial shapes of traveling waves such as sources, sinks, and other complex, heterogeneous attractors (**Figures 2**, **3**), in addition to the plane or spiral waves.

Consider **anisotropic coupling**. This can generate plane waves if, for example, the coupling is stronger along one direction than the other and if we have “**circular anisotropy**”, then we can generate rotating waves (Bachschmid-Romano et al., 2026). For example, previously, using anisotropic coupling, we were able to generate constant phase-lag waves in the lamprey central pattern generator (Cohen et al., 1992). However, as these are connectivity strengths, which are anatomically fixed, it’s not possible for them to rapidly vary from moment-to-moment or trial-to-trial or epoch-to-epoch on the fly. For example, using this model it’s not feasible to generate a source wave in one stable epoch and then a complex wave or a spiral wave in the next stable epoch. Therefore, even though this mechanism can produce only one type of wave which is again constrained by the anatomical connectivity, it cannot capture the rapid changes in the shapes of the complex spatial patterns of traveling waves that exist in our dataset.

#### Neural mass models and spiking neuron models

We have also considered other mechanistic models such as **neural mass models** and **spiking neuron models**. For example, a combination of neural mass modeling and cortico-thalamic eigenmodes apparently can generate large scale rotating waves (Gabay et al., 2018). In typical implementations of these models, the local neural circuitry consists of populations of E and I cells tuned, so that they can generate spontaneous oscillations and then they are spatially coupled via distance-dependent coupling. If they are tuned to be excitable (that is, there is a stable equilibrium, but sufficient perturbations can drive large trajectories through phase space), then if the coupling is local enough, persistent spirals can be generated by this mechanism (Huang et al., 2004). However, like our spiral waves in the flattened network, these spirals also arise from a topological phase singularity (Winfree, 1991). Therefore, the excitable spirals generated from these models and the phase-model spirals are essentially arising from the same mechanism in discrete media (Paullet & Ermentrout, 1994). A key point with the neural mass and spiking networks models is that if the coupling between local oscillation is weak, then one can obtain phase-models directly from them. However, the critical issue is that the connection between a specific neural mass model or a spiking neuron model and the phase-reduced version of it cannot be deduced except by applying the phase reduction procedure to the specific model. However, how the shape of the *H* function varies with the parameters in the neural mass model is near impossible to determine. Therefore, it was not feasible to apply these models to our working memory dataset.

#### Conclusion

Since our data is depicted as a phase map of the ECoG electrodes, phase oscillator models make the fewest assumptions and have the fewest free parameters compared to all the other alternative models considered here and they also allow for a direct comparison across trials and behavior. In contrast to, say, trying to fit the filtered LFP data at each electrode to some neural mass or spiking model would require changing many parameters even if we assume that each local oscillating circuit has the same parameters except their applied DC drive (which determines their frequency) and even then, it would be a very difficult task to produce the correct timing differences between the ECoG electrodes.

### Statistical analysis

#### (i) Identifying significant modes in CICA

To estimate the number of significant modes in the CICA output for each cluster, we compared the variance explained for each mode with the theoretical variance threshold 100/*n*, where *n* is the number of electrodes in an ECoG grid. This theoretical variance corresponds to the variance of each mode if the total variance (100%) is equally distributed among all modes. Because of the spatial structure of traveling waves, some modes will explain more variance compared to the other modes in the empirical data (**Figure 2**). We additionally shuffled the electrodes in each cluster and recalculated the variance distribution across modes and confirmed that the shuffled variance for all modes converged to the theoretical variance threshold of 100/*n*.

#### (ii) Multivariate analysis of variance (MANOVA)

To identify how traveling wave patterns related to behavior (**Figure 3**), we directly compared the weights estimated from the CICA procedure above between encoding, maintenance, and retrieval periods for each mode using multivariate analysis of variance (MANOVA). Please note that for comparison across behaviors, the stable epochs are concatenated across all task conditions and passed as inputs to the CICA and the corresponding activation functions related to the different behaviors are estimated from CICA. MANOVA was used because the weights were complex.

We statistically distinguished weights corresponding to different behavioral states using the following model: *Real + Imag ∼ States*, where *Real* and *Imag* are the real and imaginary parts of the weights respectively and *States* are encoding, maintenance, and retrieval. We used this model for each mode and applied FDR-corrections for multiple comparisons (*p* < 0.05) across all modes and subjects. Statistical significance from the MANOVA would indicate that traveling waves shift their direction and/or strength to form distinct directional patterns which can distinguish different behavioral states. We designated an oscillation cluster to be significant if at least one of the modes from the CICA for that cluster showed statistical significance in MANOVA. Moreover, we also repeated our MANOVA analysis by shuffling the labels of the behavioral states and built a histogram from these shuffles (f-statistics) against which we compared the empirical data. This analysis revealed that all our MANOVA results were still statistically significant (all *p’s* < 0.05).

We adopted similar MANOVA analysis to identify how the center of the hidden spirals related to behavior. We statistically distinguished the centers of the hidden spirals corresponding to different behavioral states using the following model: *Y + Z ∼ States*, where *Y* and *Z* are the Y-coordinates (anterior-posterior direction) and Z-coordinates (superior-inferior direction) respectively and *States* are encoding, maintenance, and retrieval. We used this model for each of the TFP and TPF oriented hidden spirals separately and applied FDR-corrections for multiple comparisons (*p* < 0.05) across all orientations and subjects. Statistical significance from the MANOVA would indicate that the centers of the hidden spirals spatially shift to distinguish different behavioral states.

#### (iii) Robustness of traveling waves at the individual epoch level

We conducted surrogate analysis to test the significance of the estimated stable epochs (see ***Identification of stable epochs*** section above) and whether the observed stable epochs are beyond chance levels. For each time-point, we shuffled the trial labels so that the temporal contiguity within a given trial is destroyed and also the electrodes, so that the spatial topography for the corresponding behavioral state is destroyed, and then ran the stable epoch analysis using identical methodology as above. In this way, we built a surrogate distribution by aggregating all time-points corresponding to these shuffled stable epochs against which we then compared the aggregated time-points from the empirical stable epochs (*p* < 0.05).

Additionally, we also carried out statistical tests based on spatial autocorrelation-preserving surrogates to ensure that the traveling waves corresponding to the different behavioral states are statistically robust. To do this, we circularly rotated the coordinates of the electrodes for each stable epoch and passed these shuffled wave patterns into our CICA approach and subjected the corresponding results to MANOVA to estimate the f-statistics for these shuffled wave patterns. Surrogates tests are powerful non-parametric tests that are adaptive to the data in consideration, and highly robust to the specific distribution of the data, therefore reducing Type I and Type II statistical errors usually found in parametric tests. F-statistics from empirical data are then compared to this histogram of f-statistics and then significance is assessed based on whether the empirical data is outside 95% (*p* < 0.05), 99% (*p* < 0.01), or 99.9% (*p* < 0.001), of the histogram. Note that this shuffling preserves the spatial autocorrelation of the wave patterns, but destroys the specific structure associated with them, therefore destroying the systematic relationship between the wave direction and task state. We then constructed the histogram of the f-statistics of these surrogates and then compared the empirically estimated f-statistics to this surrogate distribution. This analysis revealed that our results were highly significant across all subjects (all *p’s* < 0.001).

#### (iv) Robustness of hidden spirals

To ensure that the hidden spirals that we detected from the stable epochs across different behavioral states are statistically robust, we similarly carried out statistical tests based on spatial autocorrelation-preserving surrogates. This analysis will show whether the hidden spirals we detected are merely due to the spatial correlations present across electrodes or they are due to the specific structure associated with the phase patterns. We circularly rotated the coordinates of the electrodes for each stable epoch and repeated our method of flattening frequencies on these shuffled phase patterns. We repeated this procedure 1000 times for each subject. We then counted how many of these shuffled phase patterns resulted in a hidden spiral. This analysis revealed that none of the shuffled phase patterns resulted in a hidden spiral in any subject, strongly suggesting that the hidden spirals that we detected are due to the specific structure associated with the phase patterns in the stable epochs rather than spatially autocorrelated noise.

#### (v) Assessing group-level significance

For assessing statistical significance for the group level results related to the correlation coefficient between the activation functions from CICA and the hidden patterns (synchrony/spiral) for the different wave types (**Figure 10D**), we used the Kruskal-Wallis test (*p* < 0.05). These results were then subsequently FDR-corrected for multiple comparisons across task conditions.

## Acknowledgements

This research was supported by an NSF CRCNS grant to J.J. and B.E.

## Code availability

The codes for this project are available at https://github.com/anupdas777/hidden_spirals/tree/main.

## Supplementary File for

### Supplementary figures

**Supplementary Figure 1:**
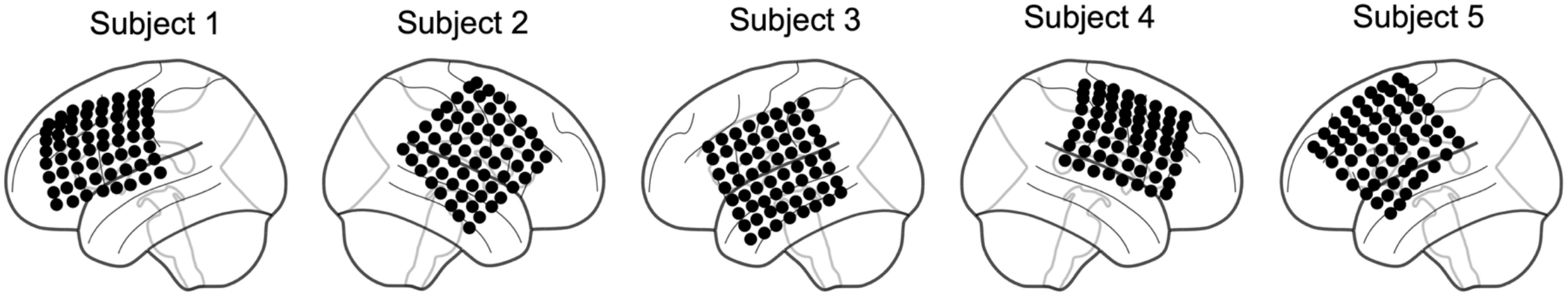
8×8 ECoG grids in the 5 subjects.

**Supplementary Figure 2:**
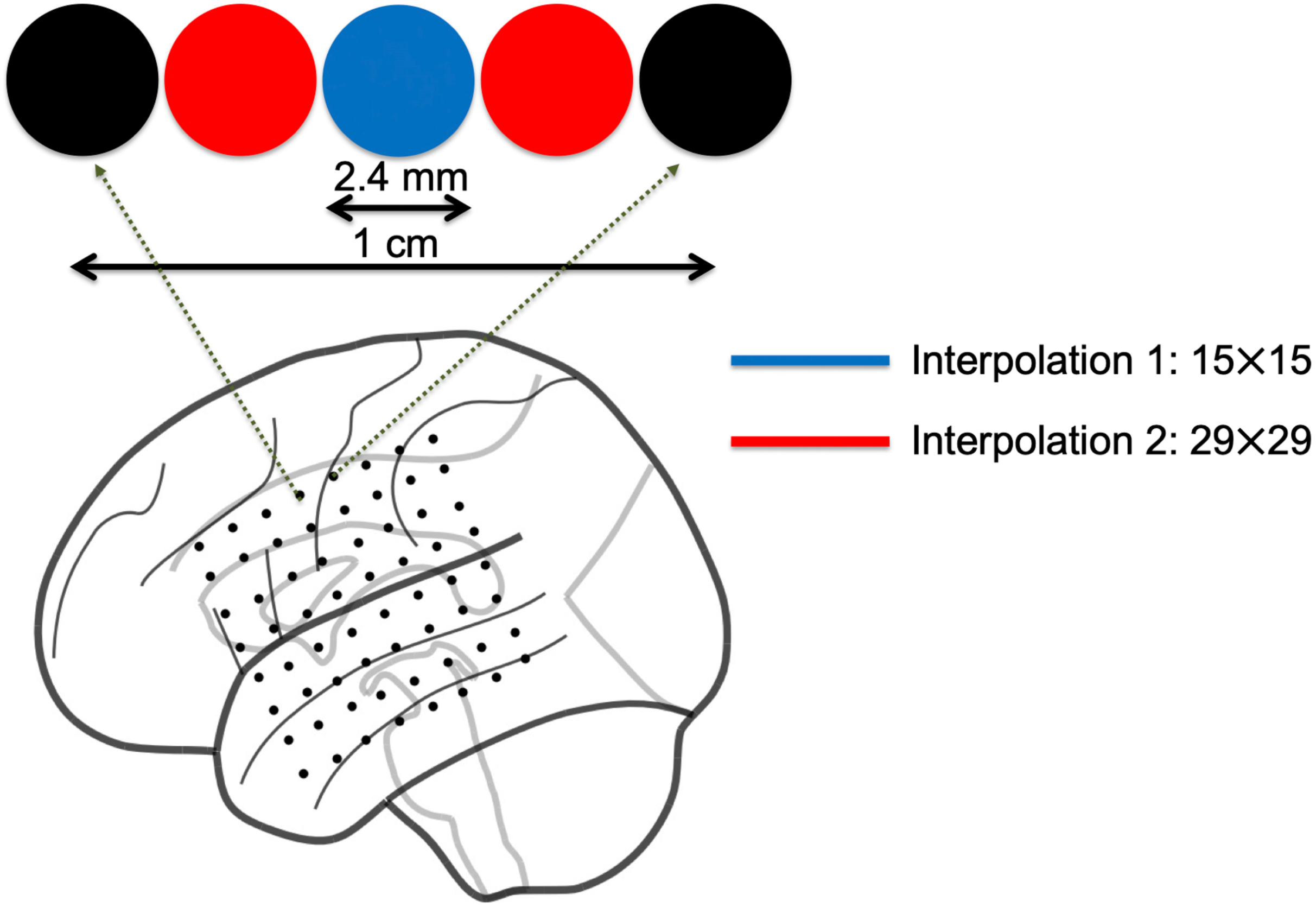
Justification for using the 29×29 phase interpolation method for hidden spiral analysis. Shown is the ECoG grid for Subject 3, with the diameter and adjacent distance of the ECoG electrodes scaled accordingly to closely match their original dimensions. As shown, the adjacent ECoG electrode distances are 10 mm, so a maximum of 3 additional shadow ECoG electrodes can be packed in between any two adjacent ECoG electrodes since the diameter of an ECoG electrode is 2.4 mm. Therefore, we first apply the first phase interpolation (blue electrode) to form the 15×15 grid, and then the second phase interpolation (red electrodes) to form the 29×29 grid.

**Supplementary Figure 3:**
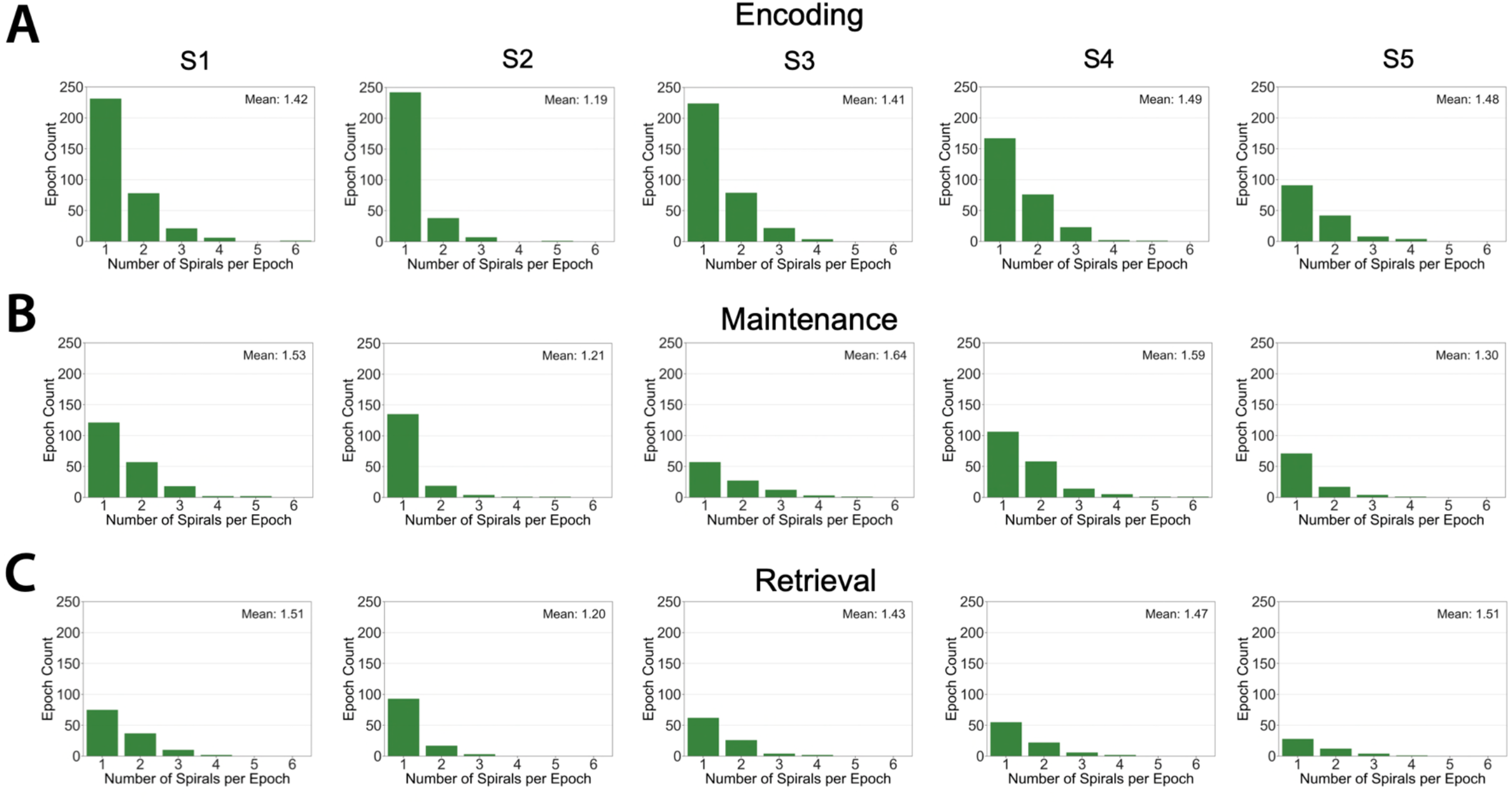
Distribution of the number of hidden spirals per stable epoch across subjects S1-S5 and across task conditions encoding, maintenance, and retrieval. Mean number of stable epochs are indicated inside each panel.

